# Nup42 safeguards heat-induced mRNAs from nuclear condensation to support chaperone synthesis

**DOI:** 10.64898/2026.02.10.705189

**Authors:** Eduardo Tassoni-Tsuchida, Angel Madero, Magda Zaoralová, Sebastian Alfonso, Brian Alford, Onn Brandman

**Affiliations:** Department of Biochemistry, Stanford University, Stanford, United States of America; Department of Biology, Stanford University, Stanford, United States of America; Department of Bioengineering, Stanford University, Stanford, United States of America

## Abstract

Cells exposed to acute stress selectively express stress-adaptive genes while repressing growth-related genes. Upon heat shock, most pre-existing mRNAs localize to translationally repressed biomolecular condensates. How heat-induced mRNAs evade condensation and remain translationally competent remains unclear. Here, we show that ribosomal protein-coding transcripts preferentially accumulate in condensates during heat shock, whereas heat-induced chaperone mRNAs are selectively excluded and preferentially translated. Using a whole-genome CRISPRi screening platform, Fractionation of Reporter-Seq (FRep-Seq), we identify the nucleoporin Nup42 as the strongest suppressor of heat-induced mRNA condensation. Loss of Nup42 triggers temperature- and transcription-dependent nuclear condensation of chaperone mRNAs, which are exported but remain translationally incompetent, leading to impaired chaperone production and thermosensitivity. Co-transcriptional mRNP packaging is a critical determinant of condensation in the absence of Nup42. Together, our findings reveal a nuclear, translation-independent layer of mRNP solubility control that enables heat shock gene expression.

## Introduction

Surviving exposure to acute environmental fluctuations requires cells to rapidly change their gene expression program. Because ribosome biogenesis and mRNA translation are among the most energetically costly processes in the cell^1,2^, these pathways are promptly repressed during stress to conserve energy^3,4^ and prevent the accumulation of misfolded or aggregation-prone proteins such as orphan ribosomal proteins (oRPs)^5–7^. Simultaneously, cells induce adaptive stress-responsive programs, most prominently the heat shock response (HSR), which upregulates chaperone expression to restore protein homeostasis (proteostasis)^8–10^. In eukaryotes, the HSR is driven by the transcription factor Hsf1, which binds heat shock elements in the promoters of heat shock protein (HSP) genes to stimulate their transcription^11–14^. A decline in proteostasis due to misregulation of stress-responsive pathways is associated with aging and neurodegeneration^15–17^. Understanding how cells manage the simultaneous challenges of preferentially expressing stress-response genes when bulk translation is repressed is a major question in the stress response field.

One proposed mechanism by which cells preferentially translate stress-induced transcripts involves the formation of biomolecular condensates, such as stress granules (SGs) and P-bodies, that form upon stress^18–20^. These condensates can sequester non-stress transcripts while excluding stress-induced mRNAs, allowing for rapid prioritization of translation ^21–25^. Consistent with this model, stress-induced mRNAs are depleted from condensates upon heat shock in yeast^22,24^, plants^26^ and human cells^27^, and similarly after glucose starvation in yeast^21^. Recent studies further suggest that the timing of transcription dictates whether a transcript condenses or is translated upon stress, with newly synthesized stress-induced mRNAs preferentially depleted from condensates^24^ and preferentially translated^24,25^. These findings suggest that processes acting co- and/or post-transcriptionally are crucial in determining the condensation status of nascent transcripts upon stress.

During transcription, nascent mRNAs are packaged into messenger ribonucleoprotein (mRNP) particles through the coordinated loading of cap-binding complex (CBC), the transcription and elongation (TREX) complex, and poly(A)-binding proteins^28^. These RNA-binding proteins (RBPs) and accessory factors are critical for proper mRNP maturation, compaction, protection from degradation and promotion of nuclear export through the nuclear pore complex (NPC)^29^. The final step of mRNP export occurs at the cytosolic interface of the NPC and relies on the catalytic activity of the DEAD-box RNA helicase Dbp5/DDX19, which removes the mRNA exporters Mex67/Mtr2 (NXT1/NXF1 in humans)^30–32^. Exposure to stress alters several aspects of mRNP processing and export^33^. Notably, heat-induced transcripts are compacted into mRNPs with distinct RBP composition, with increased Yra1 (ALYREF in humans) association and higher compaction^34^, and bypass nuclear quality control for immediate export^35^. Additionally, heat shock induces the formation of transcriptional condensates of Hsf1, Mediator, and RNA polymerase II (Pol II), reorganizing HSP loci in three-dimensional space and enhancing cellular fitness^36–38^. Yet despite the extensive characterization of the HSR regulation at the transcriptional level, the post-transcriptional mechanisms that enable heat-induced transcripts to escape condensation remain largely unknown.

Here, we combine biochemical fractionation, transcriptomics, and a genomic technique we call Fractionation of ReporterSeq (FRep-Seq) to characterize heat-specific mRNA condensation and identify its genetic regulators. We show that ribosomal protein transcripts are enriched in condensates during heat shock, while heat-induced chaperone mRNAs are depleted and preferentially translated. By performing FRep-Seq under basal and distinct time points of heat shock, we uncovered both known and previously uncharacterized regulators of transcript condensation and identified the nucleoporin Nup42 as the strongest suppressor of heat-induced mRNA condensation at 42°C. Our data suggest this function is specific to Nup42, as perturbations to other components of the same mRNA export pathway do not phenocopy the abnormal condensation of *nup42Δ* cells. In the absence of Nup42, newly synthesized chaperone mRNAs condense in the nucleus yet are still exported to the cytosol as translationally-incompetent mRNPs. Formation of nuclear condensates in *nup42*Δ cells correlates with the loss of chaperone protein synthesis, which can be alleviated by perturbations to factors involved in mRNP packaging. Together, these results demonstrate a previously unrecognized, translation-independent mechanism of nuclear mRNP condensation controlled by Nup42 that enables translation of chaperone mRNAs during heat shock to support restoration of proteostasis.

## Results

### Heat-induced transcripts are depleted from RNA condensates and are preferentially translated during heat shock

To study the transcriptomic composition of biomolecular condensates, we established a fractionation-based approach to enrich for biomolecular condensates. Our protocol did not include an immunoprecipitation step for a known condensing protein and thus did not enrich for specific types of condensates (e.g. SGs or P-bodies) (Figure 1A). Heat shock increased the levels of the yeast SG marker Pab1 (poly(A)-binding protein 1) in pellet fractions (Figure 1B and 1C), consistent with enrichment of condensates.

**Figure 1.**
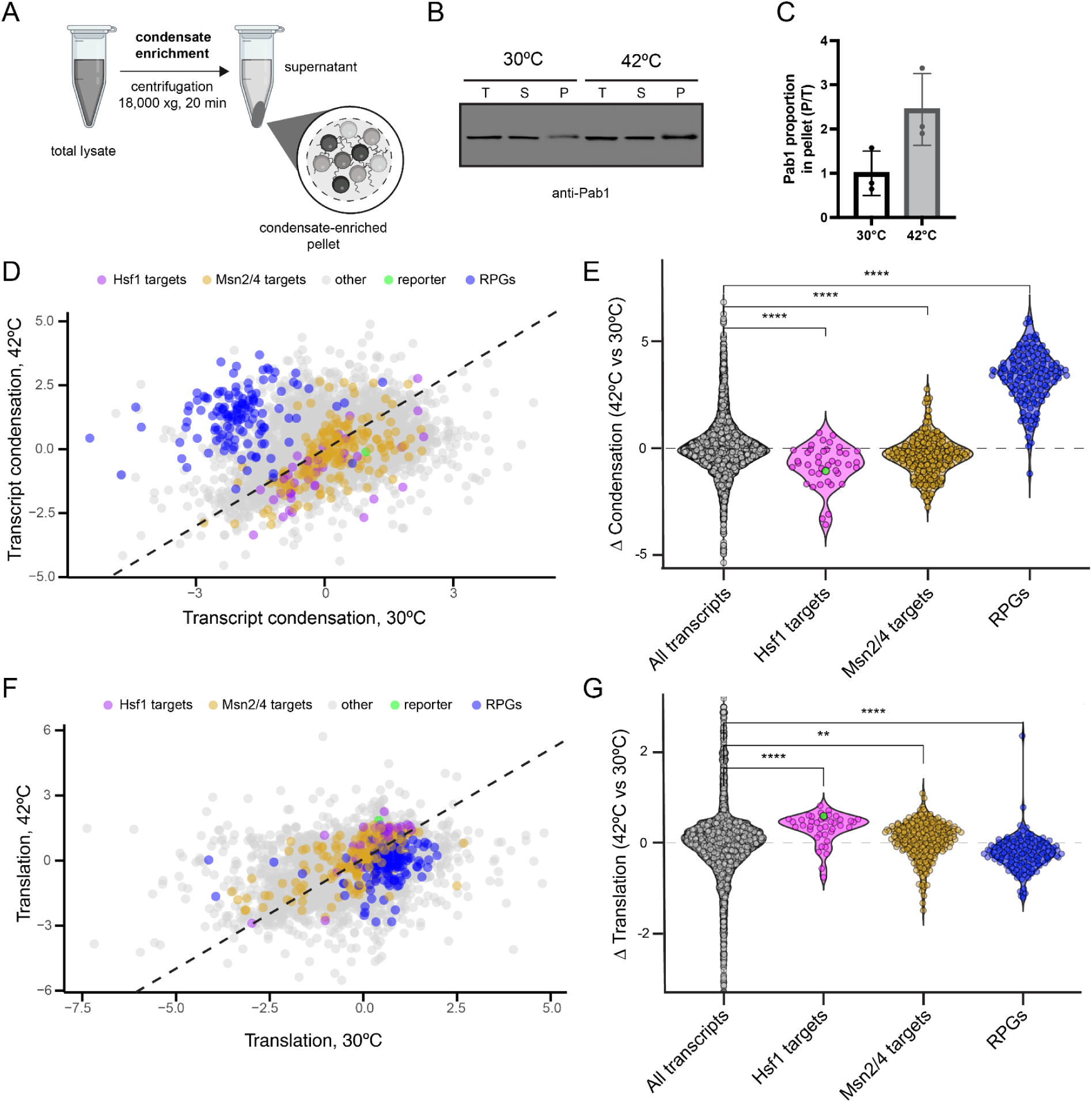
Hsf1 targets are depleted from RNA condensates and preferentially translated upon heat shock. A. Biochemical fractionation approach for enrichment of biomolecular condensates. B. Pab1 levels were monitored in the total (T), condensate-depleted supernatant (S), and condensate-enriched pellet (P) fractions at 30°C and 42°C (30 minutes) by western blotting. C. Increased condensation of Pab1 upon 42°C heat shock, measured by Pab1 signal intensity in the pellet relative to total (P/T) fractions. D. Scatterplot of length-normalized transcript condensation at 30°C and 42°C. List of Hsf1 targets (magenta) from Pincus et al., 2018^42^. Reporter (green): synthetic Hsf1 reporter encoding for GFP. Yellow represents genes regulated by Msn2/4 from Solis et al., 2016^43^. RPGs (blue): ribosomal protein genes. The dashed line represents a perfect correlation with slope = 1. E. Violin plots of changes in condensation at 42°C versus 30°C for Hsf1 targets, Msn2/4 targets, and RPGs. Green dot represents the synthetic Hsf1 reporter. Wilcoxon test with Bonferroni correction, ** (p < 0.01), *** (p < 0.001), **** (p < 0.0001). F. Scatterplot of transcript association with ribosomes, isolated by sucrose cushion, at 30°C and 42°C. The dashed line represents a perfect correlation with slope = 1. G. Violin plot of changes in ribosomal association for transcripts at 42°C versus 30°C. Wilcoxon test with Bonferroni correction, ** (p < 0.01), *** (p < 0.001), **** (p < 0.0001).

To uncover the transcriptome of condensates, we performed RNA sequencing (RNA-seq) of condensate-depleted supernatant, condensate-enriched pellet, and total fractions from a yeast strain containing an integrated Hsf1-responsive synthetic reporter coding for GFP^39,40^, under basal and 42°C heat shock conditions (Figure S1A). RNA-seq of heat shock cells showed a robust activation of both the Hsf1-driven HSR and the Msn2/Msn4-driven general stress response (GSR), along with a strong repression of ribosomal protein encoding genes (RPGs) (Figure S1B). The synthetic Hsf1-driven reporter was robustly upregulated upon heat shock, behaving similarly to other heat-induced chaperone transcripts (Figure S1B).

To quantify RNA enrichment in condensates, we used the DESeq2 algorithm^41^ to compare the distribution of reads of the condensate-enriched pellet fraction with the overall transcript abundance in the total fraction in both untreated and heat shock conditions. Longer transcripts associated with condensates regardless of stress (Figure S1C and S1D). We therefore assessed condensation after normalizing for transcript length by comparing condensation propensity of transcripts with similar sizes (see Methods). Transcript groups exhibited distinct condensation propensities, independent of transcript length but dependent on heat shock, and were replicated with an alternative quantification method (SedSeqQuant)^24^ (Figure 1D, 1E, S1C-S1E). RPG transcripts were depleted from condensates under basal conditions, but represented one of the main transcript groups enriched in heat-induced condensates (Figure 1D and 1E). In contrast, heat-induced transcripts, especially those under control of Hsf1 and including the Hsf1-responsive reporter, showed a strong depletion from heat-induced condensates (Figure 1D and 1E).

We next explored how transcript condensation correlates with translation efficiency. To enrich for translated transcripts, we isolated ribosomes from the condensate-depleted supernatant fractions using a sucrose cushion. Heat shock decreased the ribosomal association of RPG transcripts and increased the ribosomal association of heat-induced transcripts (including the Hsf1 reporter) (Figure 1F and 1G). Our data thus reveal opposing behaviours of RPG and heat-induced transcripts. RPG transcripts undergo condensation and are poorly translated upon heat shock, whereas heat-induced transcripts, especially those under control of Hsf1, evade condensation and are preferentially translated, consistent with recent findings^24^.

### FRep-Seq: a screening technology to identify modulators of RNA condensation

To investigate how heat-induced transcripts are able to escape condensation, we built upon our previously published CRISPR interference (CRISPRi) screening platform, ReporterSeq^40^, to develop a technique called FRep-Seq (Fractionation of ReporterSeq). FRep-Seq includes an additional cellular fractionation step to enrich for biomolecular condensates, enabling identification of perturbations that alter the condensation state of a reporter mRNA.

ReporterSeq investigates the genetic regulation of transcription through the usage of synthetic reporters that respond to specific transcription factors in combination with genome-wide CRISPRi perturbations. Each reporter mRNA carries a unique 3’ UTR barcode which can be used to link reporter mRNA expression to a specific gRNA, allowing genome-wide screening using a pooled plasmid library. Deep-sequencing of barcodes provides a quantitative readout of reporter abundance across all perturbations and identifies genetic modulators of transcription factor activity. In our previous study, transcription of the reporter mRNA was driven by Hsf1^40^. By adding a fractionation step to enrich for condensates and sequencing barcode counts in the condensate and total fractions, we repurposed FRep-Seq to identify biological regulators that affect Hsf1 reporter mRNA condensation (Figure 2A). Because FRep-Seq assesses condensation of the same reporter transcript across many genetic perturbations, intrinsic mRNA features (e.g. length, cis-regulatory sequences) are largely controlled for, and most of the hits are likely to reflect extrinsic regulation of condensation, although indirect effects to reporter intrinsic features may still occur.

**Figure 2.**
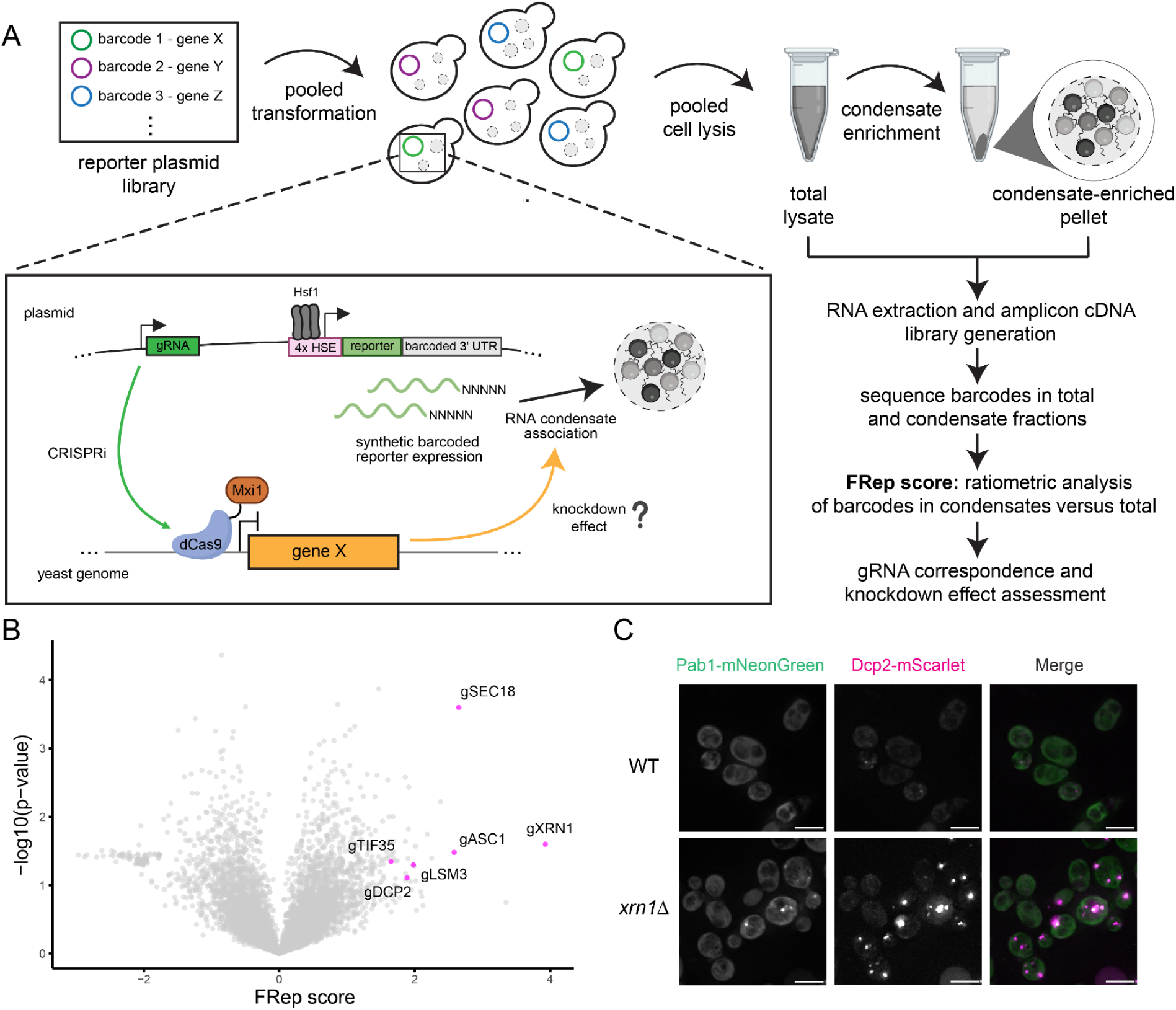
FRep-Seq uncovers genome-wide modulators of reporter mRNA condensation at 30°C steady-state. A. Fractionation of Reporter-Seq (FRep-Seq) approach: a reporter plasmid library containing gRNAs for every gene in the yeast genome linked to unique barcode sequences in the 3’ UTR of a synthetic Hsf1-responsive reporter is transformed into a pool of yeast cells already expressing constitutive dCas-Mxi1 from a multicopy plasmid. Gene knockdown effect in reporter condensation is assessed by ratiometric analysis of sequenced barcodes in condensate versus total fractions, after pooled cell lysate, condensate enrichment through fractionation, RNA extraction, reporter cDNA library generation and deep-sequencing. B. Volcano plot of FRep-Seq results of reporter basal condensation at 30°C. Every dot is a genetic knockdown, with the prefix “g” being used to facilitate the interpretation of CRISPRi perturbations. C. Fluorescence microscopy of condensate markers, Pab1-mNeonGreen for SGs, and Dcp2-mScarlet for P-bodies, in WT and *xrn1Δ* cells. Scale bar = 5 µm.

Since transcripts can condense even in the absence of stress^24^ (Figure 1D, Figure S1C), we first performed FRep-Seq in cells growing at 30°C to identify basal modulators of reporter mRNA condensation. Similar to our previous study, we used the neighborhood normalized score (NNS), a Z-score-based metric to calculate a FRep-score. Briefly, each barcode receives a score that quantifies the association of its RNA within the condensate fraction by normalizing the condensate-associated RNA to total RNA ratio against barcodes with comparable expression levels (see Methods). The sign of the score, positive or negative, indicates enrichment or depletion in condensates, respectively, while the magnitude of the score indicates the number of standard deviations from the mean. We then quantified the effect of each genetic knockdown in affecting basal reporter condensation in four biological replicates and used a one-sample t-test to quantify the reproducibility of the hits identified (Figure 2B). Knockdown of *XRN1* was the strongest hit in the positive direction (FRep score = 3.9), suggesting that loss of Xrn1 increases reporter mRNA condensation (Figure 2B). Xrn1 is a 5’-3’ exonuclease that degrades mRNAs and which deletion leads to increased P-body formation^44^. Unlike knockdown of the Hsp40 chaperone and HSR regulator *SIS1*, which robustly increases overall reporter abundance^40^ but does not cause major changes in mRNA condensation, knockdown of *XRN1* increases reporter mRNA condensation without major changes in reporter expression (Figure S2A, S2B). As expected, knockout of *XRN1* led to increased P-body formation as measured by Dcp2 foci in unstressed cells (Figure 2C), validating that FRep-Seq can identify genetic modulators of condensates. Other positive hits included additional components of P-bodies, including Dcp2, Lsm3, which have been previously shown to increase P-body formation upon impairment^44^ (Figure 2B). Knockdown of the translation initiation factor eIF3g encoded by *TIF35* as well as the ribosome-associated scaffolding protein Asc1 (RACK1 in humans) were also strong hits that increased reporter condensation (Figure 2B).

To test if hits were specific to Hsf1-driven messages, we conducted an orthogonal FRep-Seq screen using a reporter that shares the same architecture as the Hsf1-driven library except for the enhancer region of the promoter, which was controlled by the transcription factor that regulates expression of proteasome and related genes, Rpn4^45,46^ (Figure S2C). Like their effect on the Hsf1-driven reporter, depletion of both *XRN1* and *ASC1* increased condensation of the Rpn4-driven reporter under steady-state conditions (Figure S2C). By contrast, perturbations in *LSM3*, *DCP2* and *TIF35* modulated condensation of the Hsf1, but not the Rpn4-responsive reporter (Figure S2C). Thus, FRep-Seq identifies both broadly acting and promoter-specific modulators of mRNA condensation.

### FRep-Seq uncovers modulators of heat-induced RNA condensation

We next applied FRep-Seq to investigate genetic modulators of reporter mRNA condensation. Since condensation is affected by stress severity and duration^19^, we performed FRep-Seq at three time points during a 42°C heat shock: 10 minutes (early), 30 minutes (mid), and 60 minutes (late), to distinguish regulators with distinct temporal effects (Figure 3A). We observed little correlation (r = 0.03) between modulators of reporter condensation in basal and 10-minute heat shock condition (Figure 3B), indicating that distinct pathways regulate condensation at steady state compared to heat stress. In contrast, a stronger correlation was observed between the early vs mid (r = 0.44, Figure S3A) and mid vs late (r = 0.57, Figure S3B) heat shock conditions, suggesting many shared heat-specific regulatory programs throughout the time course.

**Figure 3.**
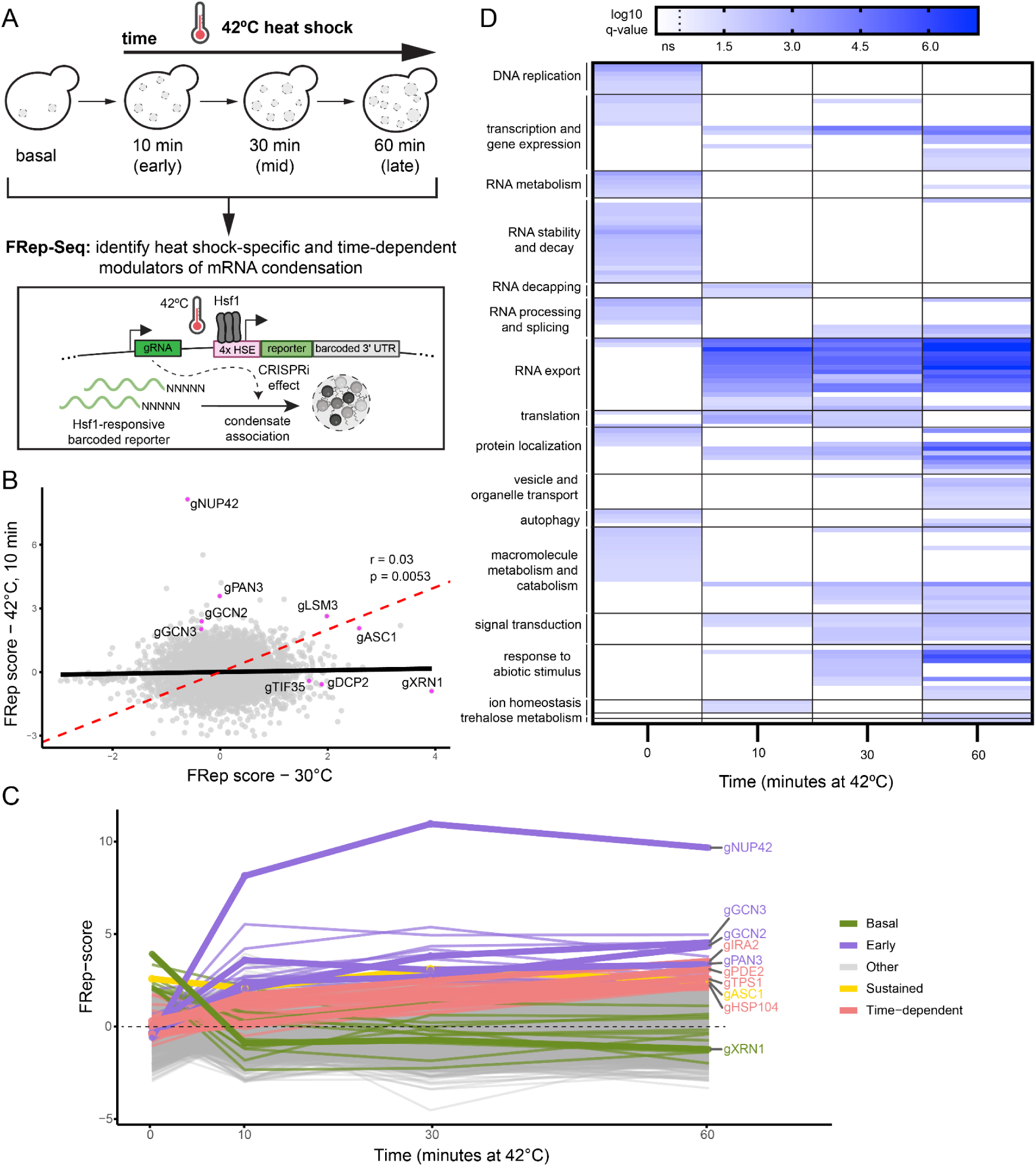
FRep-Seq reveals modulators of heat-induced reporter mRNA condensation during heat shock. A. Schematics of FRep-Seq employment to study mRNA condensation upon heat shock. B. Scatterplot of FRep scores of Hsf1-responsive reporter condensation at 30°C steady-state versus the earliest time point of heat shock at 42°C (10 minutes). C. XY plot of time (minutes at 42°C) versus FRep-score for the synthetic Hsf1-responsive reporter. Green lines represent the “Basal” group. Purple lines represent the “Early” responders group. Red lines represent the “Time-dependent” responders group. The yellow line represents the “Sustained” responder group. Gray lines represent the “Other” group, which includes genes that were not included in the previous categories. D. Gene set enrichment analysis (GSEA) of biological processes enriched among genes whose perturbations increase condensation of the Hsf1 reporter measured by FRep-Seq across all heat shock time points.

To better resolve stress- and time-dependent effects, we grouped genes based on when their knockdown increased reporter condensation in four distinct groups: basal regulators, which act only at steady state; sustained regulators, which affect condensation at both steady state and heat shock conditions; time-dependent regulators, which show additive effects over the course of heat stress; and early regulators, which affect condensation at the earliest time point of heat shock (Figure 3C). Despite their effects observed in basal conditions (Figure 2B), *XRN1* and *ASC1* fell into distinct groups: *XRN1* as a basal regulator, whereas *ASC1* showed sustained effects across both basal and heat shock conditions (Figure 3B, 3C). Time-dependent regulators included *TPS1* and *HSP104*, consistent with their respective known roles in trehalose metabolism and disaggregation, both associated with condensate dynamics under stress^47–49^. Among the early regulators, we identified *NUP42*, *GCN2*, and *GCN3*. Notably, *NUP42* emerged as the strongest hit of the screen under heat shock (Figure 3C). Nup42 is a nucleoporin with characterized roles in mRNA export from the nucleus specifically under heat shock and ethanol stress, but not under basal conditions^50–52^.

To further test the specificity of hits, we applied FRep-Seq using the orthogonal Rpn4-responsive reporter at the 30-minute heat shock time point (Figure S3C). Modulators of condensation of the two reporters showed a moderate correlation (r = 0.32), indicating some shared regulatory mechanisms (Figure S3C). Notably, *NUP42* was the strongest conserved hit between the two screens, followed by *ASC1*, whereas *GCN2* and *GCN3* preferentially affected condensation of the Hsf1-responsive reporter. These results highlight Nup42 as a robust suppressor of heat-induced mRNA condensation across multiple stress-responsive transcripts.

Beyond analysis of individual genes, we conducted gene set enrichment analysis to identify biological processes or cellular components whose perturbation increased condensation association under steady state or heat shock conditions (Figure 3D, S3D). Consistent with the weak overlap between individual genes at steady state and early heat shock (Figure 3B), enriched gene ontologies showed little overlap between these conditions (Figure 3D). Biological processes related to RNA export from the nucleus and nuclear pore-associated cellular components were the strongest and most conserved enriched terms across all heat shock time points (Figure 3D, S3D). Together, these results indicate that nuclear mRNA export pathways play a central role in regulating heat-induced mRNA condensation and identify Nup42 as a crucial suppressor of condensation of heat-induced transcripts.

### Nup42 suppresses heat-induced mRNA condensation and sustains chaperone translation

Nup42 is a conserved, non-essential nucleoporin that localizes to the cytoplasmic filaments of the nuclear pore complex (NPC), where it acts together with Gle1, the DEAD-box RNA helicase Dbp5/DDX19, and the nucleoporin Nup159 to remove the export factors Mex67-Mtr2 (NXT1/NXF1 in humans) at the last step of mRNP export from the nucleus, resulting in translationally-competent mRNPs (Figure 4A)^30–32,53,54^. Consistent with its classification as a heat-specific early regulator of condensation (Figure 3C), depletion or deletion of *NUP42* strongly increased heat-induced condensation of both the Hsf1-responsive reporter and an endogenous Msn2/4-driven chaperone gene, *HSP12*, with minimal effects to basal condensation (Figure 4B, S4A, S4B).

**Figure 4.**
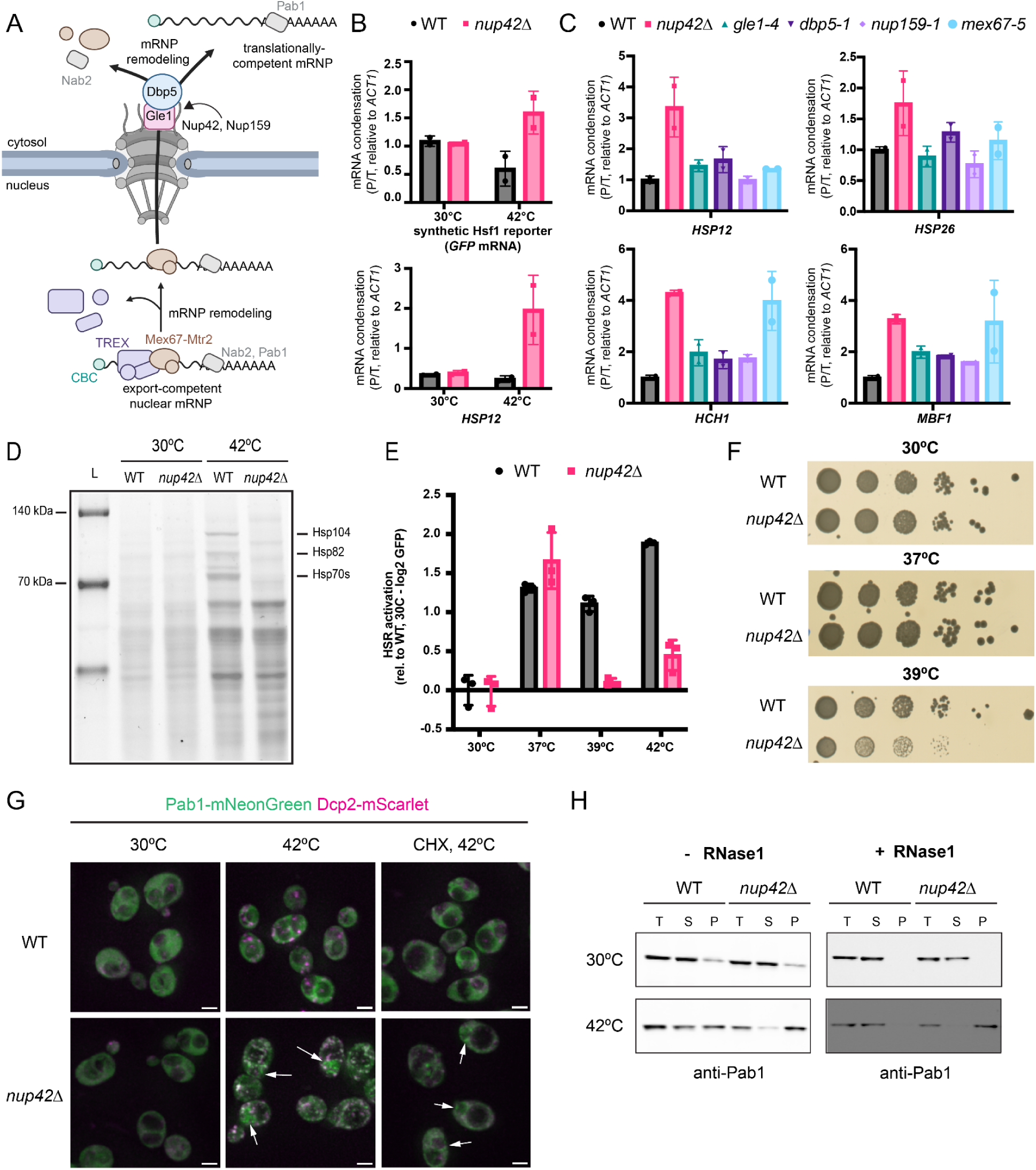
Nup42 is required to suppress heat-induced mRNA condensation and sustain chaperone protein production upon heat shock. A. Schematics of canonical Nup42 function at the nuclear pore in promoting mRNP export. B. qPCR analysis of Hsf1-responsive *GFP* reporter and *HSP12* condensation in WT and *nup42Δ* cells. Condensation measured as -ΔΔCt of pellet versus total fractions, relative to *ACT1*. N = 2 biological replicates. C. qPCR analysis of heat-induced mRNA condensation in WT, *nup42Δ, gle1-4*, *dbp5-1*, *nup159-1*, and *mex67-5* cells. Condensation measured as -ΔΔCt of pellet versus total fractions, relative to *ACT1*, then normalized to WT condensation. N = 2 biological replicates. D. Nascent protein synthesis analysis with HPG-Alexa Fluor at 30°C and 42°C heat shock for 30 minutes. E. HSR activation by GFP fluorescence measurements by flow cytometry. Heat shock was performed at the indicated temperatures for 30 minutes. CHX was added after 30 minutes of heat shock at 42°C and fluorescence was measured 30 minutes after recovery at 30°C to allow for GFP maturation. Data normalized to the mean fluorescence levels in WT at 30°C. N = 3 biological replicates. F. Spotting assays of WT and *nup42Δ* cells grown in SCD plates for 2-3 days at the indicated temperatures. Representative data of 3 biological replicates. G. Fluorescence microscopy of Pab1-mNeonGreen and Dcp2-mScarlet localization in WT and *nup42*Δ cells at 30°C and 42°C, 30 minutes. When indicated, 100 µg/mL of cycloheximide (CHX) was added to cells 10 minutes prior to 42°C heat shock. White arrows indicate Pab1 puncta that do not colocalize with Dcp2. Scale bar = 2 µm. H. Western blot of Pab1 levels in total (T), condensate-depleted supernatant (S), and condensate-enriched pellet (P) fractions. Samples in the right panel were treated with 0.3U/µL RNase1 at room temperature for 30 minutes, after cell lysis and before fractionation.

To test whether abnormal heat-induced mRNA condensation arises from a general disruption of the last step of mRNP export from the nucleus or was specific to Nup42, we investigated the effects of perturbing *GLE1*, *DBP5*, *NUP159* and *MEX67*. Depletion of these genes in the FRep-Seq screen had minimal effects in reporter condensation in both basal and heat shock conditions (Figure S4B). To ensure complete loss of function of these essential genes, we performed experiments with strains harboring thermosensitive alleles of Gle1 (*gle1-4*), Dbp5 (*dbp5-1*), Nup159 (*nup159-1*), and Mex67 (*mex67-5*), which completely block mRNP export at non-permissive temperatures.^55–58^ Although these perturbations caused mild increases in condensation of a subset of endogenous heat-induced transcripts regulated by Hsf1 (*HSP26*, *HCH1*, *MBF1*) or Msn2/4 (*HSP12*) during heat shock, the magnitude of condensation was consistently greater in *nup42*Δ cells for nearly all transcripts examined, typically by ∼1.5-2.5 fold (Figure 4C). Exceptions were *HCH1* and *MBF1*, which displayed comparable condensation in *nup42*Δ and *mex67-5* strains (Figure 4C). Under basal conditions, perturbations to these export factors had only minor effects on transcript condensation (Figure S4C). Together, these results suggest that Nup42 plays a unique role in suppressing heat-induced transcript condensation that is not explained by disruption of canonical mRNP export at the NPC.

We next investigated the functional consequences of heat-induced mRNA condensation in *nup42*Δ cells. Consistent with previous studies, bulk nascent protein synthesis analysis revealed a loss of chaperone protein production in *nup42*Δ cells during heat shock (Figure 4D)^50–52^. Using GFP fluorescence of the Hsf1-responsive reporter as quantitative readout, we observed that HSR activation was impaired at temperatures above 39°C, but unaltered at 30°C and 37°C (Figure 4E). This temperature threshold for Nup42-dependent chaperone protein synthesis correlates with the onset of thermosensitivity in *nup42*Δ cells (Figure 4F), suggesting that facilitating the expression of chaperones in heat shock is a critical function of Nup42.

To better understand the abnormal condensation observed in *nup42*Δ cells, we investigated the behavior of canonical condensate protein markers by monitoring P-body and SG formation with Dcp2-mScarlet and Pab1-mNeonGreen, respectively. Heat shock led to increased formation of P-bodies and this was further exacerbated in *nup42*Δ cells (Figure 4G). Strikingly, while only ∼10% of heat shocked wild-type cells contained Pab1-positive stress granules, nearly all heat shocked *nup42*Δ cells (∼97%) contained Pab1 condensates (Figure S4D), with a subpopulation of condensates not colocalizing with P-bodies and instead accumulating in a subcellular region that resembled the nucleus (Figure 4G). Pre-treatment with cycloheximide (CHX) disrupted the formation of heat-induced P-bodies in both wild-type and *nup42*Δ cells, and of SGs in heat-shocked wild-type cells (Figure 4G, S4D). Yet Pab1 condensates were still present in ∼64% of *nup42*Δ cells in this condition, mainly present at subcellular regions that did not colocalize with Dcp2 prior to CHX treatment (Figure 4G, S4D), suggesting that this subpopulation of Pab1 condensates is distinct from canonical SGs. Abnormal Pab1 condensation in heat shocked *nup42*Δ cells was also observed by biochemical fractionation (Figure 4H, S4E). Heat-induced Pab1 condensates were more resistant to RNase1 treatment in *nup42*Δ cells when compared to WT, indicating densely packaged mRNPs that are less accessible to nuclease degradation (Figure 4H).

Together, these results highlight a unique function for Nup42 in suppressing heat-induced mRNP condensation which is not shared among other factors acting on the same mRNP export pathway. In the absence of Nup42, during heat shock newly synthesized transcripts associate with translation-incompetent condensates and this correlates with reduced thermosensitivity.

### Nup42 suppresses nuclear condensation of newly synthesized transcripts during heat shock

The abnormal accumulation of heat-induced cycloheximide-resistant Pab1 condensates in *nup42*Δ cells prompted us to investigate their subcellular localization. We tagged the transmembrane ring protein of the NPC, Ndc1, with mScarlet to use as a nuclear membrane marker. Upon heat shock, ∼80% of *nup42*Δ cells exhibited Pab1 condensates that colocalized with Ndc1 (Figure 5A, 5B), with some cells exhibiting nucleoplasmic Pab1 puncta (Figure 5A). Notably, nuclear and NPC-localized foci were still observed in ∼55% of *nup42*Δ heat shocked cells pre-treated with cycloheximide (Figure 5A, 5B), suggesting that condensate formation does not rely on ongoing translation and is compositionally distinct from canonical cytosolic SGs.

**Figure 5.**
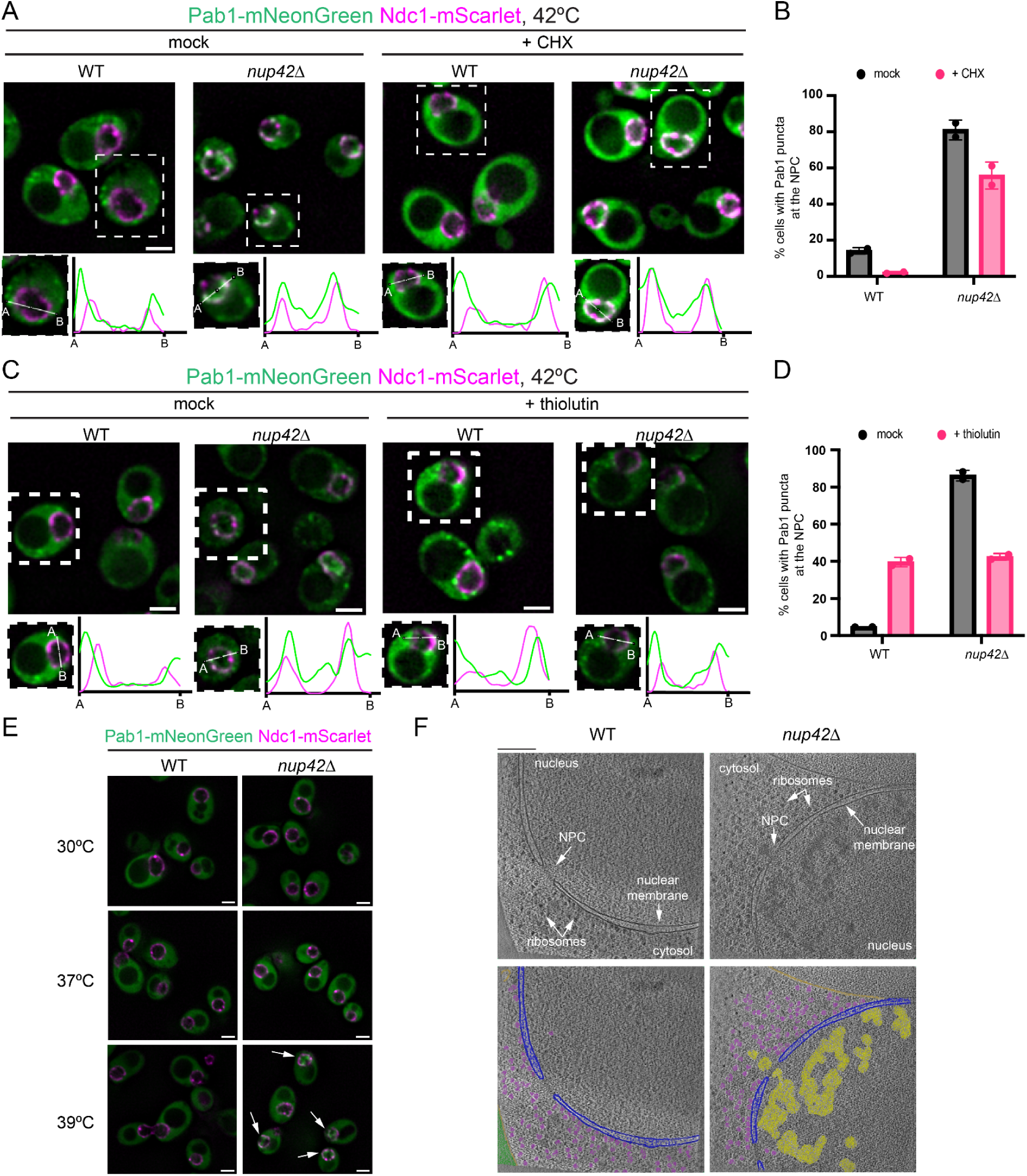
Nup42 suppresses condensation of heat-induced transcripts in the nucleus. A. Representative fluorescence microscopy Z-slice images of Pab1-mNeonGreen localization with the nuclear pore protein marker, Ndc1-mScarlet in WT and *nup42*Δ cells upon 42°C heat shock for 30 minutes. When indicated, cells were treated with 100 µg/mL of cycloheximide (CHX) 10 minutes prior to heat shock. Fluorescence intensity plots are shown for an individual cell of each condition. B. Quantification of Pab1-mNeonGreen puncta colocalization with the Ndc1-mScarlet marker as a proxy for NPC association, from panel A. N = 2 biological replicates, with at least 100 cells quantified per replicate. C. Representative fluorescence microscopy Z-slice images of Pab1-mNeonGreen localization with the nuclear pore protein marker, Ndc1-mScarlet in WT and *nup42*Δ cells upon 42°C heat shock for 30 minutes. When indicated, cells were treated with 10 µg/mL of thiolutin 10 minutes prior to heat shock. Fluorescence intensity plots are shown for an individual cell of each condition. D. Quantification of Pab1-mNeonGreen puncta colocalization with the Ndc1-mScarlet marker as a proxy for NPC association, from panel C. N = 2 biological replicates, with at least 100 cells quantified per replicate. E. Fluorescence microscopy of Pab1-mNeonGreen Ndc1-mScarlet WT and *nup42Δ* cells at the indicated temperatures (30 minutes heat shock duration). White arrows indicate nuclear Pab1 condensates. Scale bar = 2 µm. F. Top: cryo-electron tomography (cryo-ET) images of WT and *nup42*Δ upon 42°C heat shock for 30 minutes. Bottom: colored images. Purple = ribosomes, yellow = nuclear mRNP condensates, blue = nuclear membrane, golden = vacuole membrane. Scaler bar = 200 nm.

We next tested whether heat-induced nuclear Pab1 condensation in the absence of Nup42 requires nascent transcription during heat shock. Cells were pre-treated with the RNA polymerase II inhibitor thiolutin prior to heat shock, and nuclear and NPC-localized Pab1 condensate accumulation was monitored. (Figure 5C). Transcription inhibition reduced the fraction of *nup42*Δ cells displaying Pab1 foci at the NPC by ∼50% (Figure 5C, 5D). While our colocalization analysis suggested that∼42% of cells remained with Pab1 condensates overlapping with the NPC during heat shock (Figure 5D), profile trace analysis revealed a change in subcellular distribution upon thiolutin treatment (Figure 5C). In heat-shocked *nup42*Δ cells, thiolutin abolished nucleoplasmic Pab1 condensates while increasing Pab1 condensates on the cytosolic side of the NPC (Figure 5C). A similar phenotype was observed in heat-shocked wild-types cells treated with thiolutin (Figure 5C, 5D). Together, these results indicate that formation of nucleoplasmic Pab1 condensates in heat-shocked *nup42*Δ cells requires nascent transcription, whereas cytosolic Pab1 condensates in close proximity to the NPC form independently of heat-induced transcription.

To understand the functional significance of the nuclear Pab1-positive condensates observed in the absence of Nup42, we tested if their onset correlated with temperature-dependent defects in chaperone translation. Indeed, formation of nuclear Pab1 condensates did not occur in *nup42*Δ upon 37°C heat shock, but became evident at 39°C (Figure 5E), consistent with the temperature threshold at which translation and thermosensitivity defects start to occur (Figure 4E, 4F). These results indicate that the decrease in thermal tolerance in *nup42*Δ cells may be a result of abnormal condensation of newly synthesized mRNAs in the nucleus and loss of chaperone synthesis.

To investigate the extent of nuclear condensation using a marker-agnostic approach, we performed cryo-electron tomography (cryo-ET) on heat-shocked wild-type and *nup42*Δ cells. Whereas wild-type cells lacked accumulation of electron-dense material in the nuclei, *nup42Δ* cells accumulated large and heterogeneous electron-dense assemblies during heat shock, primarily in the nucleoplasm (Figure 5F). Such assemblies were also present in the vicinity of the NPC at both nucleoplasmic and cytoplasmic interfaces (Figure 5F), suggesting that mRNPs might still engage with the NPC for export from the nucleus.

Collectively, these results show that loss of Nup42 leads to aberrant nucleoplasmic Pab1 condensation during heat shock that depends on nascent transcription (Figure 5C, 5D). These nuclear condensates form independently of translation and emerge at temperatures that correlate with growth defects in *nup42*Δ cells (Figure 5A, 4F). Together, our data argue that nucleoplasmic condensate formation precedes defects in chaperone production at elevated temperatures in *nup42*Δ cells, thereby contributing to reduced thermotolerance.

### Loss of Nup42 leads to abnormal mRNP export from the nucleus during heat shock

The presence of mRNP condensates in close proximity to the NPC in heat-shocked *nup42*Δ cells motivated us to directly investigate heat-induced mRNA export from the nucleus. Previous studies suggested that Nup42 is essential for the export of heat shock mRNAs, such as the heat-induced chaperone *SSA4*^50–52,59^. However, these observations lacked single-molecule resolution and could not definitively determine whether export of heat-induced transcripts was completely blocked in the absence of Nup42. To address this, we tracked single molecules of the heat-induced Hsp70 chaperone *SSA4* mRNA using the MS2-MCP system^60^ in a Ndc1-mScarlet strain. While no prominent foci of *SSA4* mRNA was observed at 30°C in both wild-type and *nup42*Δ cells, *SSA4* transcripts were predominantly cytosolic in wild-type cells at all tested heat shock temperatures (Figure 6A). We observed a temperature-dependent accumulation of *SSA4* mRNA in the nucleus within a single bright nuclear foci in *nup42Δ* cells at 39°C and 42°C, but not at 37°C (Figure 6A, Figure S5A), which is consistent with previous findings^51,52,59^. Yet in stark contrast to previous reports, we detected cytosolic *SSA4* mRNA molecules in *nup42Δ* cells at both 39°C and 42°C (Figure 6A). Quantification of *SSA4* mRNA export revealed a ∼50% reduction in cytosolic *SSA4* mRNA foci in *nup42Δ* cells at 42°C compared to wild-type (Figure 6B). These results suggest that although heat-induced mRNA export is impaired in the absence of Nup42, it is not completely blocked at temperatures above 39°C.

**Figure 6.**
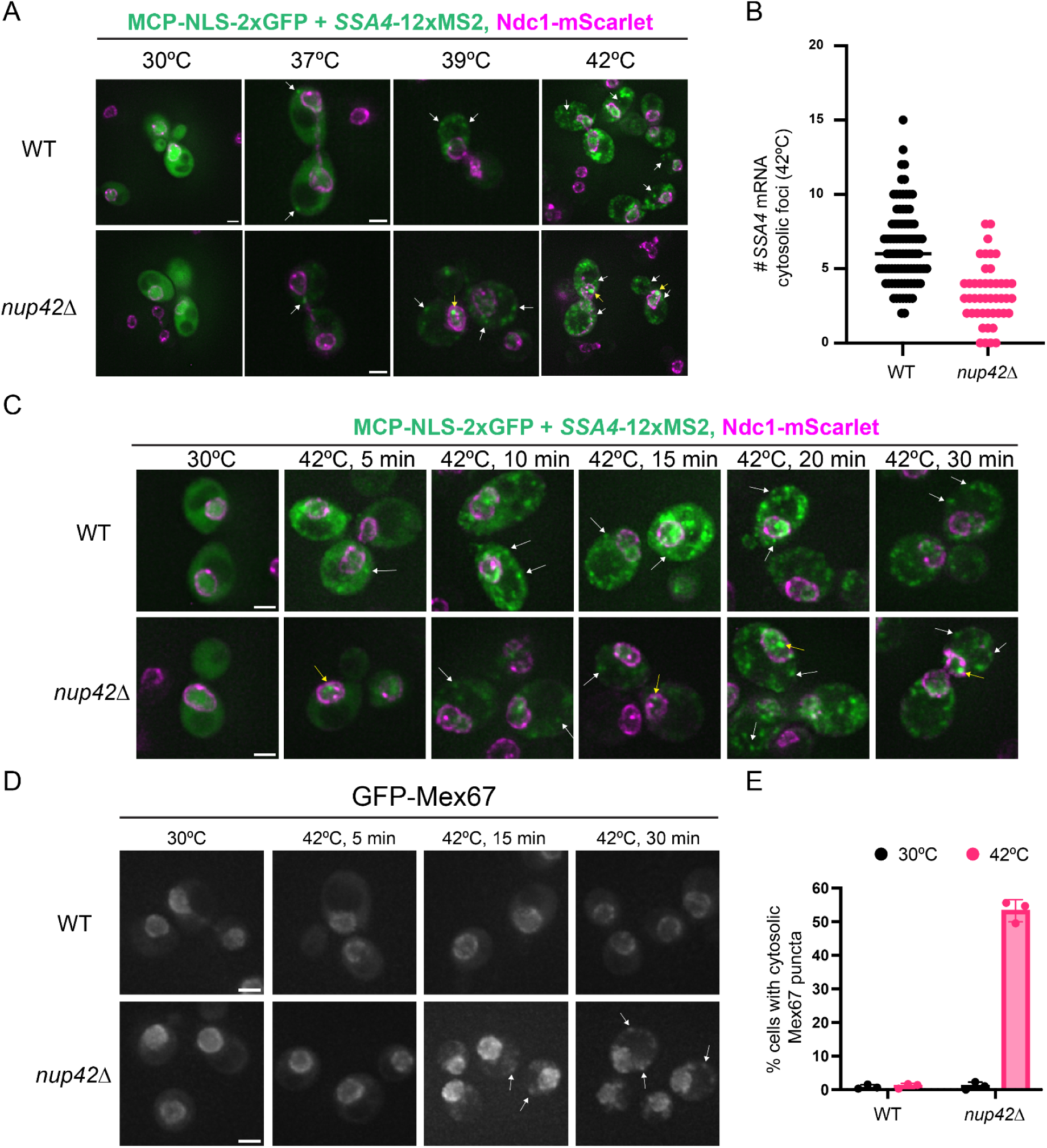
Abnormal mRNP export in *nup42Δ* cells during heat shock. A. Representative Z-slice images of single molecule *SSA4* mRNA imaging by MCP-NLS-2xGFP localization in cells with the Ndc1-mScarlet marker at the indicated temperatures. Heat shock was performed for 30 minutes. White arrows indicate cytosolic *SSA4* mRNA molecules. Yellow arrows indicate nuclear *SSA4* mRNA molecules. Scale bar = 2 µm. B. Quantification of number of cytosolic *SSA4* mRNA molecules from WT and *nup42Δ* cells at 42°C, 30 minutes shown in panel A. N = 45-90 individual cells. C. Representative Z-slice images of single molecule *SSA4* mRNA imaging by MCP-NLS-2xGFP localization in WT and *nup42Δ* cells with the Ndc1-mScarlet marker at 30°C and various timepoints of 42°C heat shock. White arrows indicate cytosolic *SSA4* mRNA molecules. Yellow arrows indicate nuclear *SSA4* mRNA molecules. Scale bar = 2 µm. D. Representative Z-stack images of GFP-Mex67 localization in WT and *nup42Δ* cells at 30°C and various timepoints of 42°C heat shock. White arrows indicate cytosolic Mex67 foci. Scale bar = 2 µm. E. Quantification of percentage of cells with cytosolic Mex67 foci from panel D. N = 3 biological replicates, with at least 100 cells quantified per replicate.

We next inspected the kinetics of mRNA export by monitoring *SSA4* mRNA localization over time during a 42°C heat shock (Figure 6C). At the 5 minute time point, cytosolic *SSA4* mRNA foci were observed in heat-shocked wild-type cells, whereas *nup42*Δ cells mainly showed nuclear foci (Figure 6C). By 10 minutes of heat shock, multiple *SSA4* mRNA molecules were detected in the cytosol of *nup42*Δ cells, while still exhibiting nuclear accumulation, suggesting delayed *SSA4* export (Figure 6C). Over the course of the 30-minute heat shock, progressively more *SSA4* mRNA molecules accumulated in the cytosol of *nup42*Δ cells (Figure 6C). However, despite the presence of cytosolic *SSA4* mRNA after 39°C or 42°C heat shock, Hsp70 protein synthesis was strongly impaired in *nup42*Δ (Figure 4D, 4E), raising the question of why exported transcripts failed to be translated.

We hypothesized that, in the absence of Nup42, heat-induced mRNAs are exported as improperly remodeled mRNPs that are still bound to nuclear export factors and remain translationally-incompetent during heat shock. To test this, we investigated whether loss of Nup42 affects displacement of the nuclear export receptor Mex67 during heat shock by monitoring GFP-Mex67 localization using fluorescence microscopy. In heat-shocked wild-type cells, Mex67 remained localized to the nucleus throughout a 30-minute heat shock at both 39°C and 42°C (Figure 6D, S5B). This phenotype was observed in *nup42*Δ at 30°C and during the first 5 minutes of heat shock at 42°C (Figure 6D). However, after 15 minutes of heat shock, *nup42*Δ cells exhibited cytosolic accumulation of Mex67 foci at both 39°C and 42°C (Figure 6D, S5B), with ∼55% of *nup42*Δ cells displaying cytosolic Mex67 foci at 42°C (Figure 6E). Remarkably, the timing of Mex67 cytosolic foci formation closely resembled the kinetics of *SSA4* mRNA export (Figure 6C), suggesting that export receptors fail to be efficiently released from nuclear mRNPs following export in the absence of Nup42. Consistent with impaired mRNP remodeling at the NPC, inspection of the canonical remodeling machinery revealed that, although Gle1 and Dbp5 remained associated with NPCs in heat-shocked wild-type cells, NPC-associated Gle1 was almost completely lost and Dbp5 was reduced by ∼70% in *nup42*Δ cells at 42°C (Figure S6A, S6B).

Taken together, our data suggest that loss of Nup42 leads to temperature-dependent nuclear mRNP condensates that engage with the NPC and are abnormally exported as translationally-incompetent mRNPs that contain Mex67, leading to defects in chaperone production and reduced thermotolerance.

### Co-transcriptional regulation of mRNP assembly affects heat-induced mRNA condensation in the absence of Nup42

To investigate what modulates abnormal condensation of heat-induced transcripts in the absence of Nup42, we performed FRep-Seq in a *nup42*Δ strain to identify conditional genetic interactions that aggravate or alleviate this phenotype, revealing factors or pathways acting upstream of or in parallel with Nup42 in modulating condensation propensity (Figure 7A). We observed a weak correlation between the FRep scores between the WT and *nup42*Δ screens (r = 0.10) (Figure 7B), indicating that loss of Nup42 substantially shapes the genetic determinants of reporter condensation.

**Figure 7.**
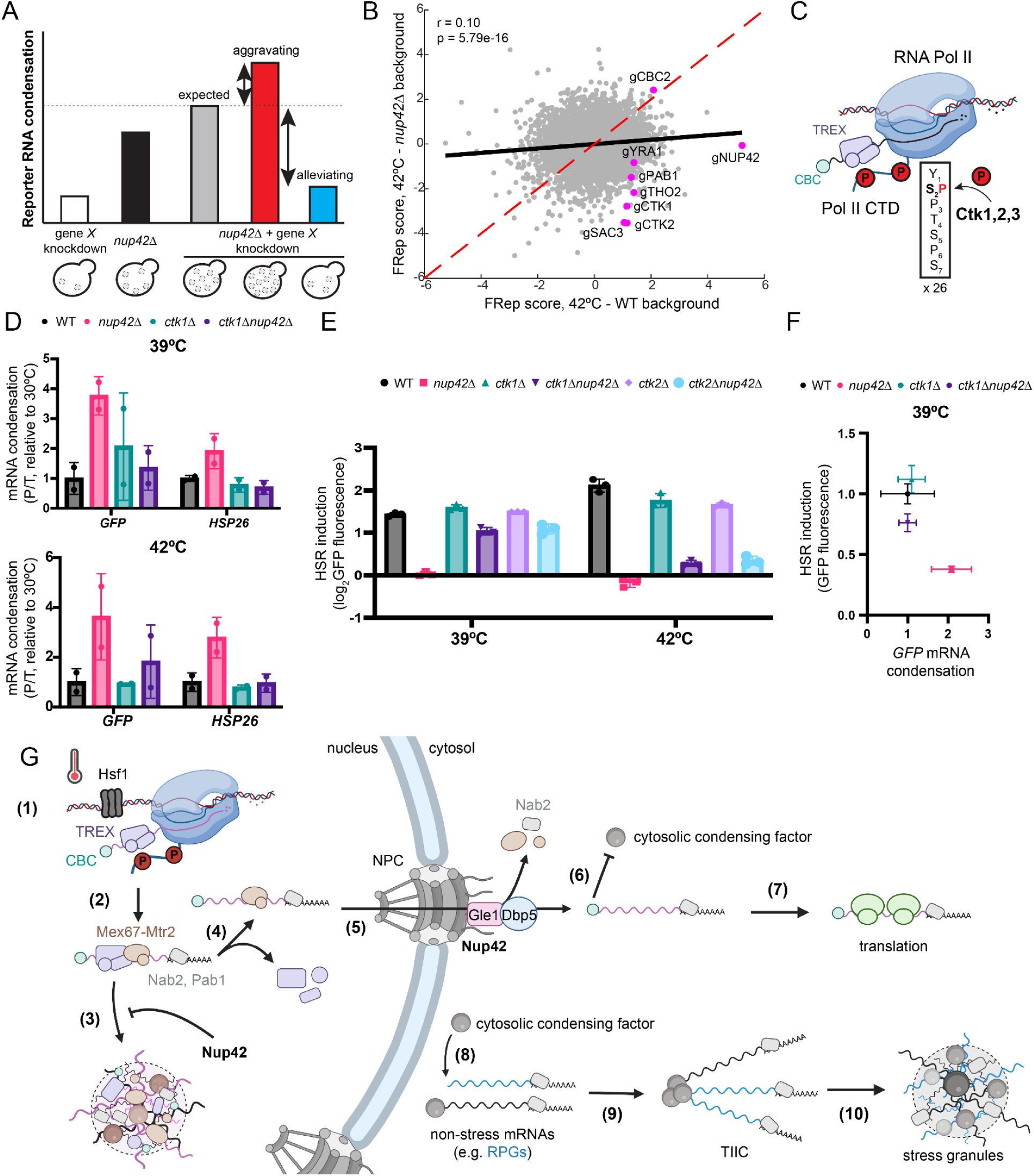
Regulation of co-transcriptional mRNP compaction modulates heat-induced mRNP condensation in the absence of Nup42. A. Schematics of genetic interaction analysis with FRep-Seq. B. Scatterplot of FRep-Seq screens conducted in WT versus *nup42*Δ background upon 42°C heat shock for 30 minutes. C. Schematics of the CTDK-I complex (Ctk1/2/3) function during transcription. D. qPCR analysis of heat-induced mRNA condensation in WT, *nup42Δ, ctk1Δ* and *ctk1Δnup42Δ* cells at the indicated temperatures. Condensation is measured as -ΔΔCt of pellet versus total fractions (P/T) upon heat shock relative to 30°C steady-state, then normalized to WT condensation for each respective transcript. N = 2 biological replicates. E. HSR activation measured by GFP fluorescence of the Hsf1-responsive reporter by flow cytometry. N = 3 biological replicates. F. Hsf1-responsive reporter transcript condensation versus protein levels upon 39°C heat shock for 30 minutes, corresponding to data shown in panels 7D and 7E. G. Unified model of transcript condensation upon heat shock. (1) Heat shock leads to Hsf1-driven transcription of heat shock genes. (2) Pol II CTD Ser2 phosphorylation by CTDK-I facilitates co-transcriptional mRNP compaction, resulting in the formation of an export-competent mature mRNP. (3) Nup42 suppresses nuclear condensation of mature mRNPs upstream of its canonical function at the NPC and (4) favors proper mRNP remodeling and NPC engagement. (5) mRNP remodeling at the cytosolic interface of the NPC by Nup42/Gle1/Dbp5 promotes dissociation of Mex67-Mtr2 export receptors and nuclear-residing RBPs. (6) Protection of 5’ cap by the cap-binding complex (CBC) inhibits cytosolic condensation of heat-induced mRNPs and (7) favors translation. (8) Concomitantly, cytosolic condensing factor(s) recognize unprotected 5’ caps of pre-existing non-stress transcripts (e.g. RPGs) causing translation initiation inhibition. (9) Global translation inhibition intensifies formation of translation-initiation-inhibited condensates (TIICs). (10) Formation of stress granules depends on TIICs and additional cytosolic condensing factors.

The strongest shared perturbation that increased reporter condensation in the two screens was the nuclear cap-binding protein *CBC2* (NBCP2), suggesting a Nup42-independent role for the cap-binding complex (CBC) in maintaining reporter solubility during heat shock (Figure 7B). A distinct subset of genetic perturbations selectively alleviated reporter condensation in the *nup42*Δ background during heat shock, with only mild effects at basal steady state (Figure 7B, S7A). Among these conditional suppressors were *PAB1*, components of the transcription-export (TREX) complex (*THO2*/THOC2, *YRA1*/ALYREF), the TREX-2 component *SAC3*/GANP, and components of the C-Terminal Domain kinase I (CTDK-I) complex, *CTK1* and *CTK2* (Figure 7B). The majority of these alleviating interactions involved factors that act either co-transcriptionally or during mRNP export. Tho2 and Yra1 function within the TREX complex in co-transcriptionally mRNP packaging^34,61^, while Sac3 is the scaffold of the TREX-2 complex which links transcription to mRNP export^62–64^. CTDK-I complex phosphorylates the YSPTSPS heptapeptide repeat of the Pol II CTD at Ser2 (analogous to CDK9/P-TEFb function in humans), promoting transcription elongation and co-transcriptional recruitment of TREX (Figure 7C)^65–68^.

Given the central role of CTD Ser2 kinases in transcription elongation and co-transcriptional mRNP packaging, we investigated the genetic interactions between *NUP42* and *CTK1*/*CTK2* in modulating heat-induced mRNA condensation. Two non-mutually exclusive models could explain the reduced condensation upon loss of Ctk1/Ctk2 in *nup42*Δ cells during heat shock. First, reduced transcriptional output due to changes in elongation rates upon perturbations to Ctk1/Ctk2 may suppress a concentration-dependent condensation regime that emerges in the absence of Nup42. Alternatively, depletion or loss of Ctk1/Ctk2 may affect co-transcriptional mRNP compaction through TREX recruitment, reducing the condensation propensity phenotype of *nup42*Δ independently of transcriptional output. To parse out these models, we measured heat-induced mRNA condensation as a function of transcript induction upon heat shock in wild-type, *nup42*Δ, *ctk1*Δ, and *ctk1*Δ*nup42*Δ cells at 39°C or 42°C. As expected, loss of Nup42 led to robust increased condensation of both the Hsf1-responsive reporter and *HSP26* mRNAs at both temperatures, whereas deletion of *CTK1* alone produced largely wild-type phenotypes under these conditions (Figure 7D). In contrast, double deletion of *NUP42* and *CTK1* substantially suppressed heat-induced transcript condensation of both mRNAs (Figure 7D). These results, together with the alleviating interactions between TREX components and *NUP42* (Figure 7B), imply that the degree of co-transcriptional mRNP compaction, rather than transcriptional output alone, are key determinants of condensation in the absence of Nup42.

We next tested whether suppression of mRNA condensation by perturbing CTDK-I complex in the absence of Nup42 was accompanied by a rescue of translation upon heat shock. As expected, loss of Nup42 severely affected translation of the Hsf1-responsive reporter at both temperatures, while deletion of *CTK1* had mild effects at 42°C (Figure 7E). Remarkably, reporter translation was restored to ∼75% of wild-type levels in both *ctk1*Δ*nup42*Δ and *ctk2*Δ*nup42*Δ at 39°C (Figure 7E, 7F), but the rescue was weaker at 42°C despite the partial rescue of reporter mRNA condensation (Figure 7E, S7B). These findings suggest that heat shock severity affects mRNA condensation status and may impose additional constraints on mRNA solubility and efficient translation. Indeed, heat-induced mRNA condensation is exacerbated at 42°C compared to 39°C when Nup42 is lost (Figure 7D).

Altogether, our results suggest that the degree and regulation of co-transcriptional mRNP compaction determines whether newly synthesized heat shock mRNPs condense and remain competent for translation in the absence of Nup42.

## Discussion

How stress-induced transcripts are selectively translated while most mRNAs are translationally repressed during acute environmental stress has largely remained a mystery. We show that heat-induced transcripts are depleted from condensates and are preferentially translated during heat shock. The suppression of heat-induced mRNA condensation requires the nucleoporin Nup42, revealing a mechanism of mRNP condensation control that operates upstream of cytosolic, translation-dependent models.

We show that transcripts can associate with condensates regardless of stress and that transcript length has minimal effects in explaining condensation propensity, which is consistent with recent observations^24^. The non-stress RPG group is among the most depleted mRNAs in condensates under basal conditions, yet dramatically increases in condensate association upon heat shock despite being relatively short (median RPG transcript length ∼1.2 kb vs. median transcript length ∼1.6 kb in yeast). Similarly, the escape of Hsf1-driven transcripts from heat-induced condensates cannot be attributed to transcript length (median ∼1.6 kb). Instead, condensation status correlates with translation repression upon heat shock, with RPG mRNAs becoming translationally repressed whereas Hsf1-driven transcripts are preferentially translated.

Glauninger et al. (2025)^24^ proposed a model in which inhibition of translation initiation through mRNA 5’ cap competition leads to the formation of translation initiation-inhibited condensates (TIICs), providing a compelling explanation for cytosolic mRNP condensation. In this framework, translation initiation repression of pre-stress transcripts triggers mRNP condensation, whereas transcription of nascent mRNAs upon stress favors translation initiation and evasion of condensation in a largely sequence-independent manner. Consistent with this model, stress-induced transcripts also preferentially escape translation repression under glucose starvation, reinforcing the hypothesis that transcriptional timing is a key determinant of mRNP translational fate^24,25^. Our findings complement this framework by showing that the condensation-fate of heat-induced transcripts is specified in the nucleus, before cytosolic translation control.

We demonstrate that upon heat shock, loss of Nup42 leads to the formation of temperature- and transcription-dependent nuclear mRNP condensates prior to export to the cytosol. Thus, heat-induced mRNP condensation competence is first determined in the nucleus rather than in the cytoplasm. We propose that the condensation-prone state of heat-induced nuclear mRNPs is co-transcriptionally established and actively suppressed by Nup42 to ensure proper mRNP export and restoration of proteostasis (Figure 7G). This model implies a previously uncharacterized anti-condensation function of Nup42 within the nucleoplasm, acting upstream of its canonical role in promoting mRNP export at the cytosolic interface of the NPC. Several observations support this framework. First, we show that heat-induced mRNP condensation is uniquely sensitive to loss of Nup42, rather than perturbing nuclear export through Gle1, Dbp5 and Nup159. Second, the genetic interactions identified in this study indicate tight cooperation between Nup42 and transcription-coupled mRNP compaction to maintain solubility of heat-induced mRNPs. Third, Nup42 localizes not only to the cytosolic face of the NPC but is also detectable within the nucleoplasm^69^. Such a model highlights previously unappreciated challenges of maintaining mRNP solubility in the nucleus upon heat shock and potentially other acute stressors.

Together with the solubilizing role of translation initiation described in the cap-competition model^24^, these observations provide a unified model for heat-induced mRNP condensation (Figure 7G). In the absence of Nup42, heat-induced mRNAs enter a condensed nuclear mRNP state that is improperly remodeled during export to the cytosol. Although the 5’ cap remains protected by the nuclear cap-binding complex (CBC), persistence of nuclear export factors such as Mex67-Mtr2 on these mRNPs may interfere with productive cytosolic remodeling and translation. This model explains both the extensive abnormal mRNP condensation observed in the nucleus and cytosol of *nup42Δ* cells and the identification of a Nup42-independent role for the CBC in suppressing cytosolic mRNP condensation during heat shock (Figure 7B, 7G). Collectively, in both the case of Nup42 and CBC, the condensation-fate of heat-induced mRNPs is specified in the nucleus, prior to cytosolic translation initiation.

What drives nuclear mRNP condensation upon heat shock? Heat shock reorganizes chromatin in three-dimensional space which promotes the formation of intergenic transcriptional condensates containing Pol II, mediator, and Hsf1^36–38^. At the same time, co-transcriptional mRNP assembly involves extensive multivalent interactions among intrinsically disordered regions (IDRs) from RBPs, nascent RNA, and the highly phosphorylated Pol II CTD. We hypothesize that these conditions increase the local heterotypic interaction density such that nascent mRNPs cross a compaction threshold and undergo abnormal condensation^70–73^. Consistent with this idea, heat-induced nuclear mRNPs exhibit altered RBP composition^34,35,74^, increased compaction^34^, and bypass nuclear quality control pathways^35^. Partial disruption of co-transcriptional mRNP packaging suppresses heat-induced nuclear mRNP condensation and alleviates translation defects of *nup42Δ* cells. Moreover, these phenotypes only emerge at higher heat shock temperatures (above 39°C) and are largely absent at 37°C. Together, these observations support a model in which heat shock severity pushes nascent mRNPs towards a condensation threshold mediated by excessive mRNP compaction, and that Nup42 buffers this transition.

Nup42 may suppress nuclear mRNP condensation by regulating the IDR interaction environment rather than binding specific mRNAs. Homotypic interactions predicted to occur among its FG-repeat region^75,76^ might limit extensive heterotypic multivalent interactions at transcription sites, reducing the effective percolation network and preventing heat-induced mRNPs from crossing an abnormal condensation threshold. In addition, Nup42 promotes engagement with the nuclear export receptors Mex67-Mtr2^53^, which may bias heat-induced mRNPs towards export-competent states and limit their dwell time at the IDR-rich environment. Subsequent stabilization of these mRNPs at the cytosolic interface of the NPC via interaction with Gle1 and stimulation of Dbp5-dependent remodeling ensures directional export and release of translation-competent mRNPs into the cytosol.^31,53,54^

Our study highlights nuclear control of mRNP solubility by Nup42 as a critical determinant of heat shock gene expression. By integrating a nuclear mechanism of mRNP condensate suppression with cytosolic models of mRNP condensation, our work provides a comprehensive framework that explains how cells achieve selective translation of stress-adaptive transcripts while enforcing widespread repression of non-essential mRNAs.

## Material and Methods

### Yeast strains and growth conditions

All the experiments performed in this study were done in the BY4741 background of *Saccharomyces cerevisiae*. For most experiments, unless otherwise indicated, cells were grown in synthetic dextrose (SD) media, with the appropriate dropout for selection, or in synthetic complete dextrose media, containing 2% w/v dextrose (Thermo Scientific), 13.4 g/L Yeast Nitrogen Base without Amino Acids (Research Products International, RPI), 0.03 g/L L-isoleucine (Sigma-Aldrich), 0.15 g/L L-valine (Sigma-Aldrich), 0.04 g/L adenine hemisulfate (Sigma-Aldrich), 0.02 g/L L-arginine (Sigma-Aldrich), 0.03 g/L L-lysine (Sigma-Aldrich), 0.05 g/L L-phenylalanine (Sigma-Aldrich), 0.2 g/L L-threonine (Sigma-Aldrich), 0.018 g/L L-methionine (Sigma-Aldrich), 0.036 g/L L-tryptophan (Sigma-Aldrich), and 0.018 g/L uracil (Sigma-Aldrich).

Cells were grown from single colonies in a temperature-controlled room at 30°C for at least 16 hours, until reaching OD_600_ = 0.2 – 0.4, before harvesting or being exposed to heat shock. For small (< 5 mL cultures), cells were centrifuged in a table-top centrifuge to pellet the cells. Heat shock was conducted by replacing supernatant media with pre-warmed media at the desired temperature (37°C, 39°C or 42°C) kept in a thermomixer containing adaptors for Falcon tubes (Thermomixer C, Eppendorf). Heat shock cells were then placed on a thermomixer with gentle shaking (800-1000 rpm) for the desired duration. Control cells were replaced with room temperature or 30°C pre-warmed media and kept in a thermomixer or in the 30°C room. For larger cultures (200 mL to 1 L), cells were harvested by vacuum filtering onto nitrocellulose membranes and pellet were scrapped with razor blades and transferred to Falcon tubes with 30-50 mL of pre-warmed media and kept in a thermomixer for the duration of the heat shock. After heat shock, cells were again vacuum filtered, and pellets were flash frozen in liquid nitrogen and stored in the -80°C for further processing.

Yeast transformations were performed using standard methods with lithium acetate (240 µL 50% PEG, 36 µL 1M lithium acetate, 10 µL 10 mg/mL single-stranded salmon sperm DNA) at 42°C for 20-40 minutes, before selection in the appropriate media. Gene deletions were obtained by replacing the endogenous ORF with either a *NATMX6*, and selected in YPD (Difco YPD Broth, BD) with 2% agar (VWR) plates with 100 µg/mL nourseothricin/clonNAT (Research Products International, RPI), or *KANMX6*, and selected in YPD agar plates with 400 µg/mL G418 sulfate (Cayman Chemical Company) through homologous recombination. Successful knockouts were confirmed by colony PCR and Sanger sequencing.

### Fractionation for condensate enrichment

Cell lysates were obtained by combining the flash frozen pellets with 1-2 mL of flash frozen lysis buffer containing 50 mM Tris HCl pH 8.0 (Thermo Fisher), 140 mM potassium acetate (Thermo Fisher), 6 mM magnesium chloride (Thermo Fisher), 0.1% NP-40 (Sigma-Aldrich), 1 mM PMSF (Thermo Fisher), Pierce EDTA-free protease cocktail inhibitor (Thermo Fisher), 0.5 mM DTT (Sigma-Aldrich), 1:5000 Antifoam B (Sigma-Aldrich), and 1 U/mL RNAseOUT (Thermo Fisher) – unless otherwise indicated. Pellets were then lysed in a freezer mill (SpexCertiPrep 6750 or 6800) filled with liquid nitrogen, at 10 Hz for 2 minutes. The lysate powder was then transferred to Falcon tubes and stored in the -80°C for further processing. Powder lysates were thawed for at least 2 hours on ice before transferred to microcentrifuge tubes for clarification at 2,000 g for 2 minutes at 4°C. Clarified supernatants were then transferred to new microcentrifuge tubes. 10% of the volume of the clarified lysate was transferred to another tube for total/unfractionated lysate analysis. The remaining 90% was centrifuged at 20,000 g for 20 minutes at 4°C to obtain a pellet enriched in biomolecular condensates. The supernatant was then transferred to a new tube that corresponds to condensate-depleted supernatant fraction. The condensate-enriched pellet was then washed with 500 – 1000 µL of freshly prepared lysis buffer and subjected to another round of centrifugation at 20,000 g for 20 minutes at 4°C. The final pellets were then resuspended in 250 µL of lysis buffer for further RNA isolation.

### Sucrose cushion and ribosomal enrichment

400 µL of the supernatant/condensate-depleted fraction from the step above was 2x diluted with lysis buffer and overlayed onto 800 µL of sucrose cushion buffer (50 mM Tris pH 7.5, 6 mM magnesium chloride, 25 mM HCl, 2M sucrose, 100 µg/mL cycloheximide, 1 U/mL RNAseOUT) in 13 x 56 mm polycarbonate thick wall tubes (Beckman Coulter, #362305). Ribosomes were obtained by ultracentrifugation at 430,000 g at 4°C for 1 hour using TLA-110 rotor (Beckman Coulter) in an Optima XL-100K ultracentrifuge (Beckman Coulter). Ribosome pellets were resuspended in 250 µL of lysis buffer for RNA isolation.

### RNA isolation

Total, supernatant, condensate-enriched and ribosome-enriched pellet fractions were combined with 10X AE Buffer (500 mM sodium acetate pH 5.2, 100 mM EDTA) in order to obtain 500 µL at 1X AE buffer concentration and transferred to screw-cap tubes. 50 µL of 10% SDS (Promega) and 500 µL acid phenol:chloroform (Thermo Fisher) was added to the tubes, vortexed, incubated to 65°C with shaking at 1400 rpm for 10 minutes in an Eppendorf Themomixer C, followed by a 5-minute incubation on ice. The solution was then centrifuged at 15,000 g, at 4°C for 15 minutes. Aqueous phase was then transferred to new tubes and 550 µL of chloroform (Fisher Chemical) was added, followed by centrifugation at 15,000 g for 5 minutes, at 4°C. The aqueous phase was transferred to new tube, then 40 µL of 3M sodium acetate pH 5.2 and 400 µL of isopropanol were added, incubated at room temperature for 10 minutes, followed by centrifugation at 15,000 g for 20 minutes, at 4°C. RNA pellets were then washed with 750 µL of 75% ethanol and centrifuged at 15,000 g for 5 minutes, at 4°C. The RNA pellet was air-dried for 3-5 minutes at room temperature before resuspension in 50 µL of nuclease-free water. RNA samples were stored in the -20°C for downstream applications.

### Western blotting

Protein samples were obtained as described above, from 0.5 – 1 mg of clarified cell lysate quantified by Pierce™ BCA (Thermo Fisher). Condensate-enriched pellets were 4X concentrated relative to the supernatant and total fractions. 4X NuPAGE™ LDS Sample Buffer (Invitrogen, #NP0008) with 5% β-mercaptoethanol (MP Biomedicals, #190242) was added to each fraction for a final 1X concentration. Samples were then boiled at 95°C for 5 minutes and 5-10 µL was loaded onto a 3-8% NuPAGE™ Tris-acetate gels (Thermo Fisher, #EA0375, #EA03755), and ran for 45-60 minutes at 150 V. Proteins were then transferred to nitrocellulose membranes (Biorad, #) Towbin buffer (25 mM Tris, 192 mM glycine, 20% w/v methanol) with wet-lab transfer apparatus (Biorad, #1704070), at 100 V for 1 hour in a 4°C cold-room. Blocking was performed for 1 hour with 5% milk in TBST buffer. Primary antibody against Pab1 (EnCor Biotechnology, #MCA-1G1) at 1:5000 in 5% milk-TBST solution was incubated overnight at 4°C. Membranes were washed three times in TBST before incubation with secondary antibody conjugated with 1:5000 IRDye 800 donkey anti-mouse IgG secondary antibody (Licor, #926-32212) in 5% milk-TBST solution, for 1 hour at room temperature. Membranes were then scanned on a Licor Odyssey CLx and bands were quantified using Fiji/ImageJ.

### RNA-Sequencing and data analysis

RNA samples from total, supernatant, condensate-enriched and ribosome-enriched pellet fractions was treated with 2 U of Turbo DNase I (ThermoFisher, #AM2238), before submitted to Bioanalyzer for quality check. RNA concentration was quantified using Qubit™ RNA HS assay kit (Invitrogen, #Q32855). Poly(A)+ mRNA was isolated using the NEBNext Ultra II RNA library prep kit for Illumina® (NEB, #E7770L) from 1,000 ng of input, using NEBNext Poly(A) mRNA magnetic isolation module (NEB, #E7490). All washes were performed with Axygen® AxyPrep MAG PCR Clean-Up beads (Corning, #MAG-PCR-CL-50). Quality check of final libraries was performed with Bioanalyzer High Sensitivity at the Stanford Protein and Nucleic Acid (PAN) facility. Libraries were quantified with Qubit™ dsDNA HS assay kit (Invitrogen, #Q32854) and equimolarly pooled together to a final concentration of 2 nM. Libraries were denatured following Illumina NextSeq protocols to a final loading concentration of 2 pM. Sequencing was performed with a NextSeq 500/550 High Output Kit v2.5 75 cycles (Illumina, #20024906), single-read, obtaining at least 5 million reads per sub-library.

The reference genome was obtained from the Saccharomyces Genome Database (SGD), version S288C_reference_genome_R64-5-1_20240529, and the CDS of the synthetic Hsf1-responsive GFP reporter was manually added to the reference genome. Reads were demultiplexed using bcl2fastq. Reads were trimmed using TrimGalore (version 0.5.0)^77^, mapped to the reference genome and gene counts were obtained with STAR (version 2.5.4b)^78^, using –quantMode. Differential gene expression was analyzed using DESeq2^41^ with design = ∼condition + fraction + condition:fraction, with 30°C total fraction serving as reference levels.

The list of genes targeted by Hsf1 and Msn2/4 was obtained from Pincus et al., 2018^42^ and Solis et al., 2016^43^. Transcript length was obtained from the most recent genome reference based on mRNA lengths with UTRs for all the coding genes.

Transcript condensation was calculated as the log2 fold change of transcript abundance in condensate-enriched pellet fractions relative to their total abundance in the total fraction, for both 30°C and 42°C conditions. To account for transcript condensation regardless of transcript length, transcripts were sorted based on length and binned into groups of 100, and a Z-score was calculated based on the behavior of each transcript within the bin of transcripts with similar lengths, similar to Glauninger et al., 2025^24^. Calculation of proportion of transcripts in the supernatant fractions was performed using the SedSeqQuant algorithm^24^, and the sedScores were compared to the condensation scores obtained from DESeq2^41^ after length normalization.

Transcript translation status was obtained through a similar approach as transcript condensation, but measuring the log2 fold change of transcript abundance associated with ribosomal fractions relative to their total abundance, for the basal and heat shock conditions.

### Reporter plasmid library construction and yeast library transformation

Hsf1-responsive reporter plasmid library was the same used in Alford & Tassoni-Tsuchida et al., 2021^40^. Rpn4-responsive reporter plasmid library was the same used in Work et al., 2021^46^. Both reporters contain a crippled CYC1 promoter sequence and encode for emerald GFP, but have distinct minimal promoters, either heat-shock elements (HSE) or proteasome-associated control elements (PACE), that respond to Hsf1 and Rpn4, respectively. Both reporter libraries contain approximately 12 guide RNAs per gene, and each guide RNA can be associated with more than one barcode to minimize PCR bias. In this study, a new set of transformations of the Hsf1-responsive reporter plasmid library in the WT and *nup42*Δ backgrounds was performed. In summary, yeast cells are transformed with the 2µ episomal pMET17-dCas9-Mxi1 / *LEU2* plasmid and grown into saturation in SD -Leu media. The cells are then diluted into a 1L YPD culture to OD_600_ = 0.1 and grown to OD_600_ = 0.4-0.6. Cells are then transformed with 1 µg of the reporter plasmid library and grown in 1 L of SD -Ura -Leu for 2-3 days, until reaching saturation. The resultant pool of cells is then concentrated, aliquoted in 1 mL with 50 OD_600_ of cells in SD -Ura -Leu media containing 25% glycerol, and stored at -80°C.

### Fractionation of ReporterSeq (FRep-Seq) and downstream analysis

Two frozen aliquots (biological replicates) of yeast containing the reporter library were thawed at room temperature and grown for 16 hours in 1 L of SD – Ura – Leu media, at 30°C. When OD_600_ = 0.4-0.5, cells were vacuum filtered and concentrated into 40 mL in SD -Ura -Leu media, and 10 mL was transferred to new Erlenmeyer flasks containing 1 L of SD -Ura -Leu media pre-warmed at 30°C overnight, in order that each flask will be subjected to a different treatment. When the diluted cultures reached OD_600_ = 0.4, cultures were vacuum filtered, harvested and flash frozen in liquid nitrogen, in the case of the untreated condition, or transferred to new Erlenmeyer flasks with 1 L of SD -Ura -Leu media pre-warmed at 42°C overnight in a shaker, for the heat shock conditions. Flasks were heat shocked for 10, 30 or 60 minutes at 42°C, vacuum filtered, and flash frozen in liquid nitrogen. Frozen pellets were lysed and processed as described in the fractionation for condensate enrichment session above. Only total and pellet fractions were subjected for further RNA isolation and cDNA amplicon library generation. RNA isolation was the same as described above. RNA samples were treated with Turbo DNase I and quality checked as described above.

Poly(A)+ mRNA selection was performed with 50 µg of input using NEXTFLEX® Poly(A) Beads 2.0 (PerkinElmer, #NOVA-512993). The resulting poly(A)+ mRNA was then reverse transcribed using Multiscribe reverse transcriptase (ThermoFisher) using a gene-specific primer that anneals to the 3’ UTR of the synthetic GFP reporter. The cDNA was purified after alkaline hydrolysis of the RNA with sodium hydroxide, followed by purification with AxyPrep MAG PCR Clean-Up beads. Barcoding PCR containing unique index sequences were added and dsDNA PCR products were gel-purified in 2% TAE agarose gels using Zymoclean Gel DNA recovery kit (Zymo Research, #D4001).

To obtain yeast plasmid DNA samples for cell/barcode count estimation, 50 mL of frozen yeast libraries was thawed and miniprepped using the Zymoprep Yeast Plasmid Miniprep Kit II (Zymo Research, #D2004). Index sequences were added by barcoding PCR and the PCR product was purified as described above.

PCR libraries were quantified using Qubit™ dsDNA HS and pooled at equimolar concentrations at a final concentration of 3 nM. Libraries were denatured and ran on a NextSeq 500/550 High Output Kit v2.5 75 cycles, single-read, as described above. Approximately 10 million reads were obtained for each sub-library.

Reads were demultiplexed and barcodes were matched to guide RNA sequences using kallisto, as described in Alford & Tassoni-Tsuchida et al., 2021^40^. The list of guide RNAs and their counts was then input into Matlab (MathWorks, R2024) for further analysis. As described in Alford & Tassoni-Tsuchida et al.,^40^, basal neighborhood normalized scores (NNS) and interaction NNS were generated by comparing pairs of conditions. Basal NNS compares differences in reporter RNA and DNA levels, serving as a proxy for transcriptional activity or abundance after gene knockdowns with CRISPR interference. Gene-stress interactions can be obtained by comparing reporter RNA levels after stress relative to unstressed RNA levels (interaction NNS). In summary, NNS allows for quantitative measurement of barcode effects while minimizing noise by using a neighborhood normalization, in which each barcode abundance score between two conditions is normalized to the distribution of other barcodes with similar counts. A positive NNS reflects genetic knockdown that increases reporter expression, while a negative NNS reflects perturbations that decrease reporter expression. The magnitude of the scores (absolute value) represents the strength of the effect in standard deviations. In this study, we named the comparison of reporter RNA levels between pellet and total fractions, FRep score, while still utilizing the same principle as the NNS for its calculation^40^. Genetic perturbations with positive FRep scores increase reporter RNA association with condensate fractions, while negative FRep scores decrease reporter condensation.

FRep scores were filtered for classification of distinct responder groups shown in Figure 3C. Basal regulators correspond to hits with FRep score > 2 at time 0 minutes of heat shock and lower than 3 at 10, 30 and 60 minutes of 42°C. Sustained regulators correspond to hits with FRep score > 2 at all time points. Early regulators correspond to hits with FRep score < 2 at 0 minutes, and > 2 at 10, 30 and 60 minutes of heat shock. Time-dependent regulators correspond to hits whose FRep score difference between time points were > 0 with a final score > 2 at 60 minutes of heat shock.

### Gene set enrichment analysis (GSEA)

FRep scores were sorted in descending order for the different conditions and heat shock time points, and the resulting gene name list was used for gene set enrichment analysis in R using the gseGO function of the clusterProfile package^79^ and reference genome org.Sc.sgd.db. A Benjamin-Hochberg correction was used, with a p-value cutoff of 0.05. GSEA was performed for biological process (BP) and cellular component (CC) gene ontology terms.

### RT-qPCR

RNA was extracted and treated with DNase as mentioned above. 5 µg of RNA was then reverse transcribed using Multiscribe reverse transcriptase (Thermo Fisher, #4311235) using random hexamers. cDNA was then mixed with Luna® Universal qPCR Master Mix (NEB, #M3003E) using gene-specific primers. Each sample-primer combination was measured in three technical replicates with a ViiA 7 qPCR machine (Thermo Fisher). The median Ct value of the three technical replicates was used in the analysis. Each qPCR data point shown in the figures represents a biological replicate. RNA induction levels after heat shock were calculated using the - ΔΔCt measurement, using *ACT1* as a reference. For transcript condensation measurements, - ΔCt values for each transcript in each condition were calculated, and these values were then normalized to the – ΔCt values of *ACT1*, unless otherwise indicated.

### Flow cytometry

Yeast cells were grown from single colonies for at least 16 hours in SCD media until reaching OD_600_ = 0.2-0.4. For heat shock treatments, 1 mL of culture was centrifuged, and pellet was resuspended in pre-warmed media at the desired temperature. 200 µL of cells were transferred to wells of a 96-well plate and plates were shaken at 1,000 rpm at the desired temperature. Since folding of the emerald GFP is disturbed at 42°C, cells were allowed to gradually recover by transferring 96-well plate from a 42°C plate shaker to a 30°C for 30 minutes, with the addition of 100 µg/mL of cycloheximide per well to cease protein synthesis during recovery. Unstressed cells were centrifuged and resuspended in room temperature media, and 200 µL were transferred to wells of a 96-well plate and kept at 30 °C plate shaker for 30-60 minutes. GFP fluorescence was then measured on a BD Accuri C6 flow cytometer with 10,000 events per sample, under the fast flow setting.

### Nascent protein synthesis measurements with HPG

Cells were grown overnight from three independent colonies in SCD medium for at least 16 hours until reaching an OD600 of 0.4–0.6. For heat-shock treatment, 1 mL of SCD–methionine medium was pre-warmed to 42 °C. Ten OD units of cells were pelleted by centrifugation and resuspended either in 30 °C SCD–methionine medium (control) or in 42 °C SCD–methionine medium supplemented with 50 µM L-homopropargylglycine (HPG; Vector Labs, Cat. #CCT-1067). Cells were incubated in 1 mL of methionine-depleted medium in Falcon tubes and shaken at 1000 rpm for 30 minutes at the indicated temperature (30 °C or 42 °C) in a Thermomixer C (Eppendorf).

Following treatment, cells were pelleted and resuspended in 1 mL of pre-chilled 70% ethanol, then fixed overnight at 4 °C. Fixed cells were washed twice with 1 mL of 3% BSA in PBS, pelleted, and incubated with 100 µL of Click-iT reaction cocktail (Invitrogen, Cat. #10428) for 30 minutes at room temperature, protected from light. Cells were pelleted and washed once with 100 µL of rinse buffer, followed by two washes with 1 mL of PBS. The final pellet was resuspended in 50 µL of 2× Laemmli buffer containing 20 mM DTT.

Cells were lysed by boiling at 95 °C for 5 minutes and centrifuged at 16,000 × g for 10 minutes at 22 °C. Lysates were diluted 1:2, and 5 µL was loaded onto a 4–20% Mini-PROTEAN TGX gel (Bio-Rad, Cat. #4561096). HPG–Alexa Fluor labeling was imaged on a Typhoon FLA9500 scanner (GE Healthcare Life Sciences). For total protein quantification, gels were stained with InstantBlue, imaged, and total lane intensity was measured as a proxy for protein loading.

### Fluorescence microscopy

Cells were grown from single colonies for at least 16 hours in appropriate media at 30°C to OD_600_ = 0.2-0.4. For heat shock treatments, 1 mL of culture was centrifuged and resuspended in pre-warmed media at the desired temperature, followed by incubation at Eppendorf Thermomixer C with 800-1,000 rpm at the desired temperature and time. After heat shock, cells were centrifuged and concentrated to 100 µL, and 50 µL was added to concanavalin-A treated wells in microscopy chambers (Cellvis, #C8-1.5H-N), with the addition of 500 µL of pre-warm media. For unstressed control cells, 1 mL of culture was centrifuged and concentrated to 100 µL and transferred to concanavalin-A treated wells, with the addition of 500 µL of room temperature media. When indicated, 100 µg/mL of cycloheximide or 10 µg/mL of thiolutin were added 10 minutes prior to the heat shock. After treatment, cells were immediately imaged using an Olympus ix83, using a 100x/1.45 oil objective, controlled with the cellSens software. 0.25 µm Z-stacks were obtained, unless otherwise indicated. Images were deconvolved using the cellSens software and Z maximum intensity projections were obtained. Images were analyzed and quantified using Fiji/ImageJ software. The number of replicates and cells counted are shown in figure legends.

### Cell culture vitrification for cryo-FIB milling

Yeast cells were grown to OD_600_ = 0.2-0.4 in SCD media. Heat shocked cells were subjected to 42°C heat shock for 30 minutes. 5 µL of cell culture was applied to a quantifoil X/X mesh grid, back-blotted for 5s and flash-frozen in liquid ethane in a Leica EM GP2 plunge freezer. The prepared grids were clipped into an autogrid system (Thermo Fisher Scientific) and stored in liquid nitrogen storage dewar prior cryo-FIB thinning.

### FIB-milling

Following vitrification, flash-frozen yeast cells on EM grids were loaded into a cryoFIB/SEM system Aquilos (Thermo Fisher Scientific) under cryogenic temperatures. Subsequently, the whole grids were coated with platinum. First layer consisted of a rough sputter coating with parameters of 10 mA, 10 Pa and 15 s using an integrated plasma coater. Then, another layer was deposited, this time of an organoplatinum (GIS) compound, approximately 400 nm of thickness, followed by another rough sputter layer (again, 10 mA, 10 Pa and 15 s). The cells were localized on grids in the SEM modality in the Maps SW (Thermo Fisher Scientific) or directly in the Aquilos xT UI (Thermo Fisher Scientific). Clusters of yeast cells (2-6 cells) were FIB-milled to approximately 170-220 nm thickness using a Gallium ion source accelerated by 30 kV in the Aquilos system. Care was taken to select well-hydrated clusters and avoid any potentially blotted areas. First, the rough cuts were performed with 1 - 0.5 nA to achieve thickness of approximately 1-2um. Then, 100 - 30 pA currents were used for thinning and polishing to the final lamella thickness. During the final steps, to monitor the thinning process and integrity of the protective platinum layer the process was monitored by imaging with an electron beam of 2 kV and 13 pA. Polished lamellae were coated with a fine sputter layer with 7 mA, 10 Pa and 8 s coating conditions.

### TEM data acquisition

Prepared lamellae were cryo-transferred to a Krios G2 (Thermo Fisher Scientific) equipped with a direct detector Falcon 4i (Thermo Fisher Scientific) and an energy filter Selectris X (Thermo Fisher Scientific). Tomographic series were collected using a Tomography 5 SW (Thermo Fisher Scientific), using multi-shot acquisition set up, dose-symmetric tilt scheme, tilt span ranging from 50 degrees and 2.5 degrees tilt step increments. Region of interest was automatically tracked and autofocusing procedure was run for each tilt on an area adjacent to a central area of interest. The pixel size was 2.461A^2^ and the defocus values ranged from 2.5 to 6 µm. The energy slit was set to 10 eV and each individual tilt dose was calibrated to receive 3.5 electron per Angstrom^2^ per projection, resulting in 140 e^2^ per entire tilt series. Data were saved in the eer format of Falcon4i detector.

### TEM data and tomographic reconstruction

For data reconstruction, tilt series in eer format were converted to tiff file format using the Relion3 implementation of SBGrid. Individual projections were motion corrected with the MotionCor2, while simultaneously dividing the dataset into even/odd frames. Frames were aligned and reconstructed with AreTomo. Reconstructed 3D volume was used for model training in CryoCare and the resulting model served as a denoising platform for the final 3D tomographic volume that was segmented using ORS Dragonfly neural networks.

### Single-molecule RNA imaging with the MS2-MCP system

The 3’ UTR of the endogenous *SSA4* gene was tagged right downstream of the stop codon with the 12xMS2V6 containing a *KANXM6* resistance cassette flanked by loxP sites from plasmid pET246-pUC57 12xMS2V6^60^, by transformation and selection in YPD + G418. Colonies that contain the cassette were confirmed by colony PCR. Positive colonies were then transformed with the pBF3038 plasmid^80^, expressing Cre recombinase under the *GAL1* promoter, were selected in SD -Leu plates for 2-3 days. A mix of colonies were pooled together and grown overnight in synthetic media containing 2% raffinose instead of dextrose as the carbon source, with dropout of leucine. The next day, cells were pelleted and resuspended in synthetic media containing 2% galactose and dropout of leucine, for expression of the Cre recombinase. Cells were kept in galactose overnight, diluted and plated into SD -Leu plates. Individual colonies were then subjected to colony PCR to monitor the loss of the *KANMX6* cassette, which was confirmed by Sanger sequencing. Colonies that lost the *KANMX6* cassette were then transformed with a modified version of the pET296-YcpLac111 CYC1p-MCP-NLS-2xyeGFP plasmid^60^, containing a *HIS3* auxotrophic marker instead of *LEU2*. Cells were grown and fluorescence images were acquired as mentioned above.

## Material availability

Strains and plasmids will be made available upon request.

## Acknowledgments

We thank Judith Frydman, Ron Kopito, Joseph Puglisi, James Ferrell, Flora Rutaganira, members of the Brandman, Kopito and Pleiner labs for helpful discussions and feedback. We thank Ben Montpetit for sharing the *dbp5-1*, *gle1-4*, *mex67-5* and *nup159-1* strains. pET246-pUC57 12xMS2V6 and pET296-YcpLac111 CYC1p-MCP-NLS-2xyeGFP were gifts from Robert Singer & Evelina Tutucci (Addgene #104390 and Addgene #104394). pBF3038 was a gift from Nancy DaSilva & Suzanne Sandmeyer (Addgene #26850). This work was supported by the NIH (R35GM153301) to O.B., Stanford DARE (Diversifying Academia, Recruiting Excellence) Fellowship to E.T.T., Stanford SGF (Graduate Fellowship in Science & Engineering) Fellowship to A.M., the Lucille P. Markey Basic Biomedical Research Fellowship and NIH 5 T32 GM007276 to B.A, and the Stanford Bio-X Undergraduate Summer Research Program to S.A.

## Author contributions

Conceptualization, E.T.T. and O.B.; methodology, E.T.T., A.M., M.Z., S.A., B.A., O.B.; investigation, E.T.T., A.M., M.Z.; formal analysis, E.T.T., A.M., M.Z., O.B.; visualization, E.T.T., A.M., M.Z., O.B.; writing - original draft, E.T.T., A.M., O.B.; writing - review and editing, E.T.T., A.M., M.Z., S.A., O.B.; funding acquisition, E.T.T., A.M., S.A., B.A., O.B.; resources, O.B.; supervision, O.B.

## Supplemental Figures

**Figure S1.**
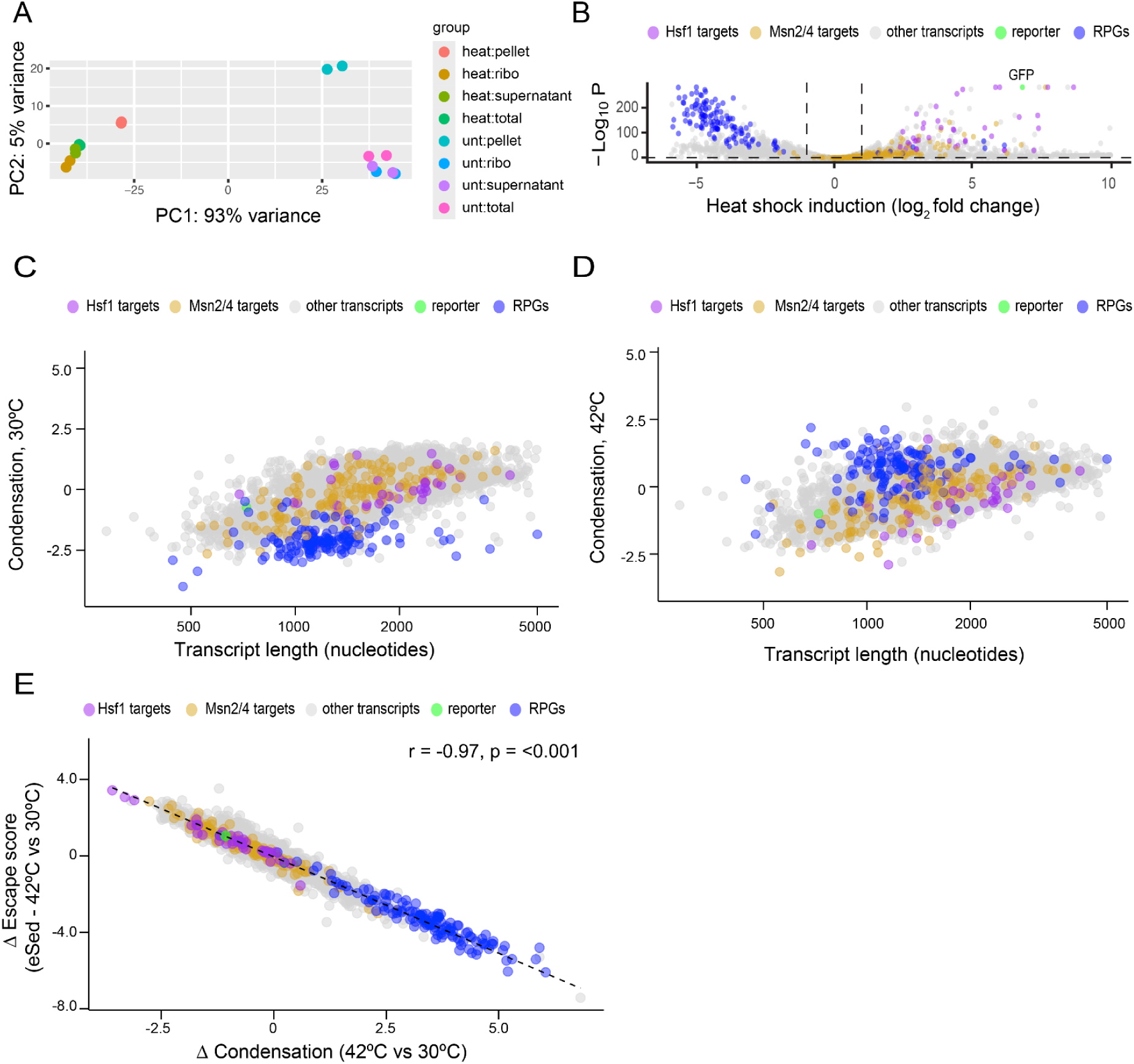
Hsf1 target and RPG transcripts condensation status is independent of length. A. PCA plot of RNA-Seq libraries by condition (unt = 30°C, heat = 42°C for 30 minutes) and fraction (pellet = condensate-enriched, ribo = ribosomal, total = unfractionated). B. Volcano plot of transcriptional changes upon 42°C heat shock for 30 minutes, showing upregulation of chaperone transcripts (Hsf1 and Msn2/4 targets), and downregulation of ribosomal protein genes (RPG). The Hsf1 synthetic reporter behaves as a canonical Hsf1 target (green). C. Transcript length versus condensation status at 30°C. D. Transcript length versus condensation status at 42°C. E. Scatterplot of length-normalized changes (42°C versus 30°C) in condensate enrichment with DESeq2 versus proportion in condensate-depleted supernatant fractions with Sed-Seq.

**Figure S2.**
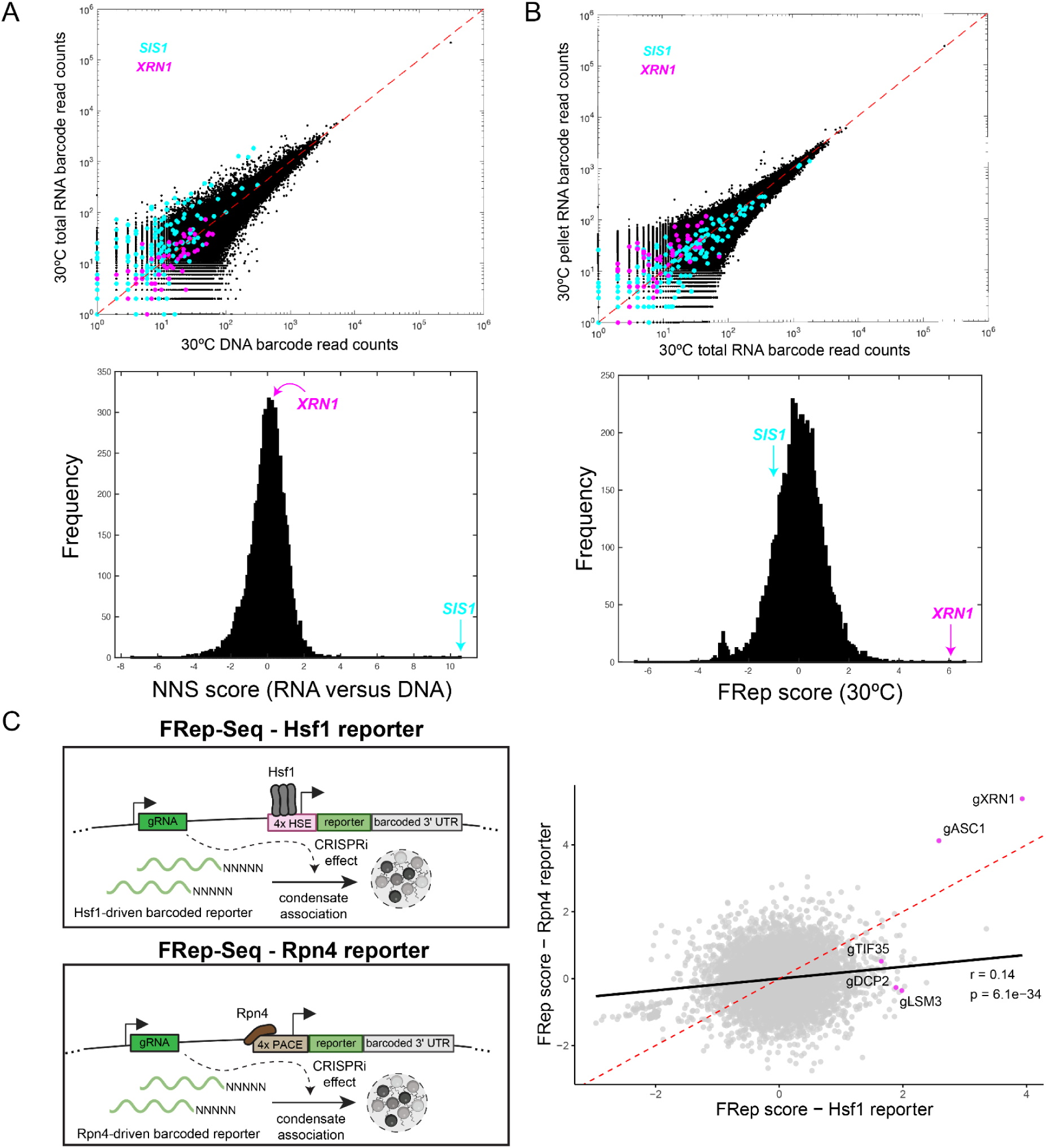
FRep-Seq identifies modulators of condensation despite of transcriptional levels. A. Top panel: scatterplot of unstressed 30°C plasmid DNA barcode reads versus RNA barcode reads from total unfractionated samples. Every dot represents a unique barcode associated with a given gene. *SIS1* represents the strongest HSR activator identified previously by ReporterSeq^40^. Bottom panel: histogram of calculated NNS scores as a proxy for HSR transcriptional activation. B. Top panel: scatterplot of unstressed 30°C RNA barcode reads from total unfractionated versus RNA from condensate-enriched fraction. Bottom panel: histogram of calculated FRep-scores as a proxy for reporter condensation. C. Left panel: schematics of Hsf1-responsive and Rpn4-responsive reporters and their usage in FRep-Seq. Right panel: scatterplot of FRep-Seq results of reporter condensation for Hsf1-dependent versus Rpn4-dependent reporters at 30°C steady-state.

**Figure S3.**
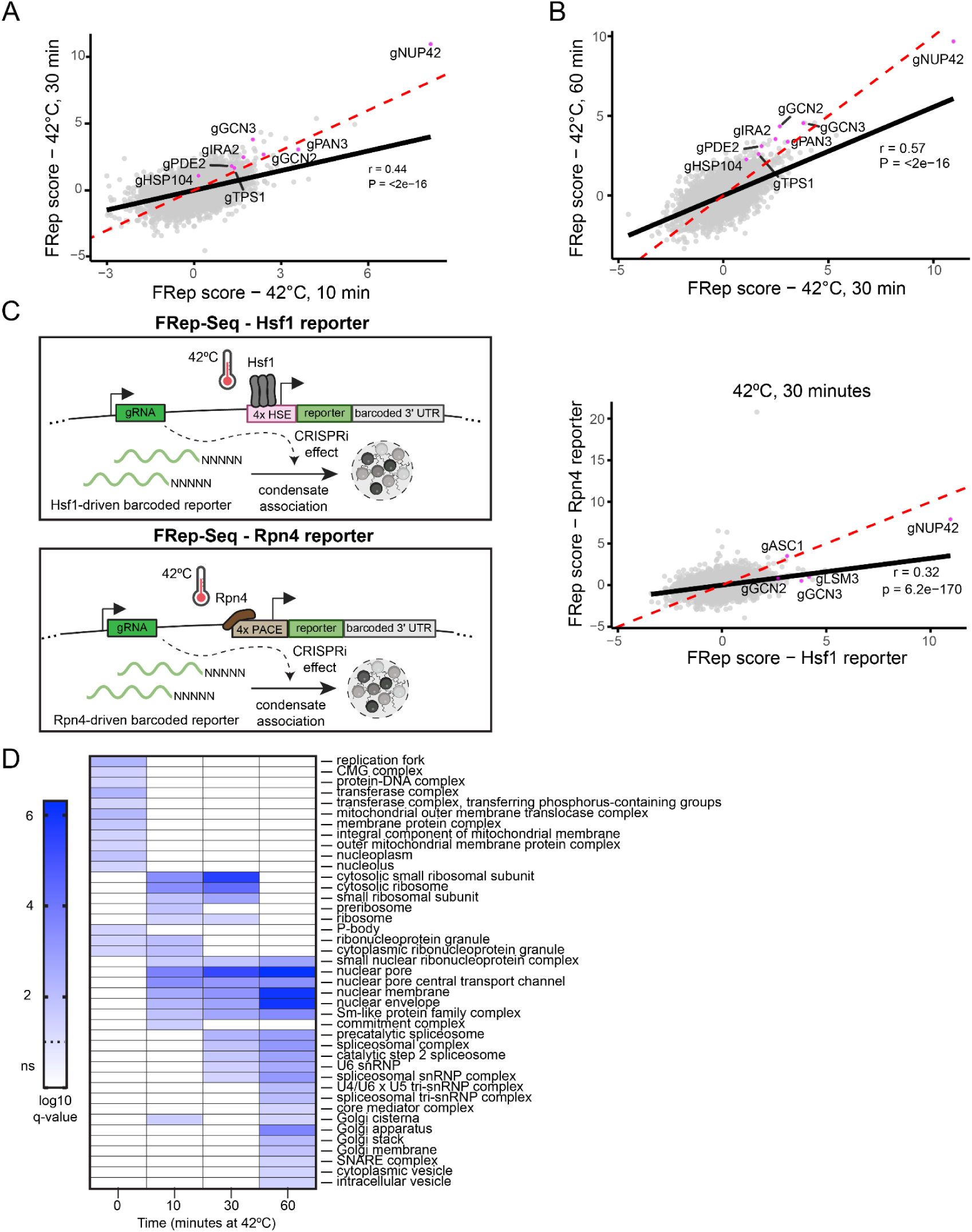
FRep-Seq reveals distinct modulators of Hsf1-reporter escape from condensation at basal and heat shock conditions. A. Scatterplot of FRep-scores for the Hsf1-responsive reporter at 10 minutes versus 30 minutes of a 42°C heat shock. B. Scatterplot of FRep-scores for the Hsf1-responsive reporter at 30 minutes versus 60 minutes of a 42°C heat shock. C. Scatterplot of FRep-scores for Hsf1-responsive versus Rpn4-responsive reporter at 42°C, 30 minutes. D. Gene set enrichment analysis (GSEA) of cellular components enriched among genes whose perturbations increase condensation of the Hsf1 reporter measured by FRep-Seq across all heat shock time points.

**Figure S4.**
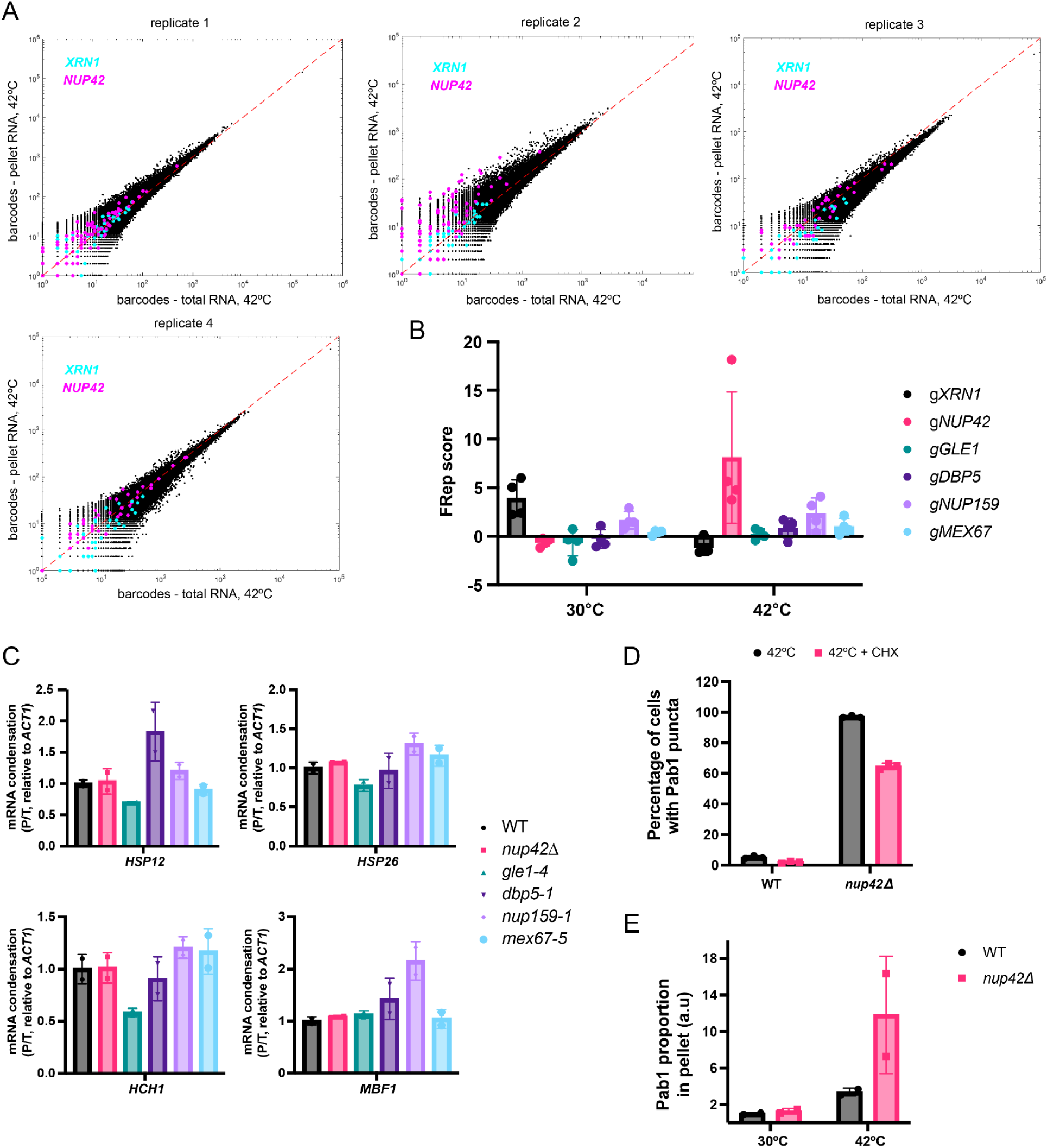
Nup42 does not affect basal condensation. A. Scatterplots of barcode reads from the total unfractionated sample versus RNA barcode reads condensate-enriched pellet upon 42°C heat shock for 30 minutes, for all four biological replicates. Replicates 3 and 4 consist of a different batch of yeast library stocks from another reporter library transformation than replicates 1 and 2. B. FRep-scores of genes shown in Figure 4B, as well as the strongest hit in the basal unstressed condition, *XRN1*. N = 4 biological replicates. C. mRNA condensation at 30°C measured as proportion in pellet relative to total abundance (P/T) for the indicated strains. N = 2 biological replicates. D. Quantification of fluorescence microscopy images shown in Figure 4G, quantifying percentage of cells with Pab1-mNeonGreen puncta. N = 3 biological replicates, with at least 100 cells quantified per replicate. E. Quantification of Pab1 proportion in pellet fractions (P/T) from lysates without RNase1 treatment shown in Figure 4H. N = 2 biological replicates.

**Figure S5.**
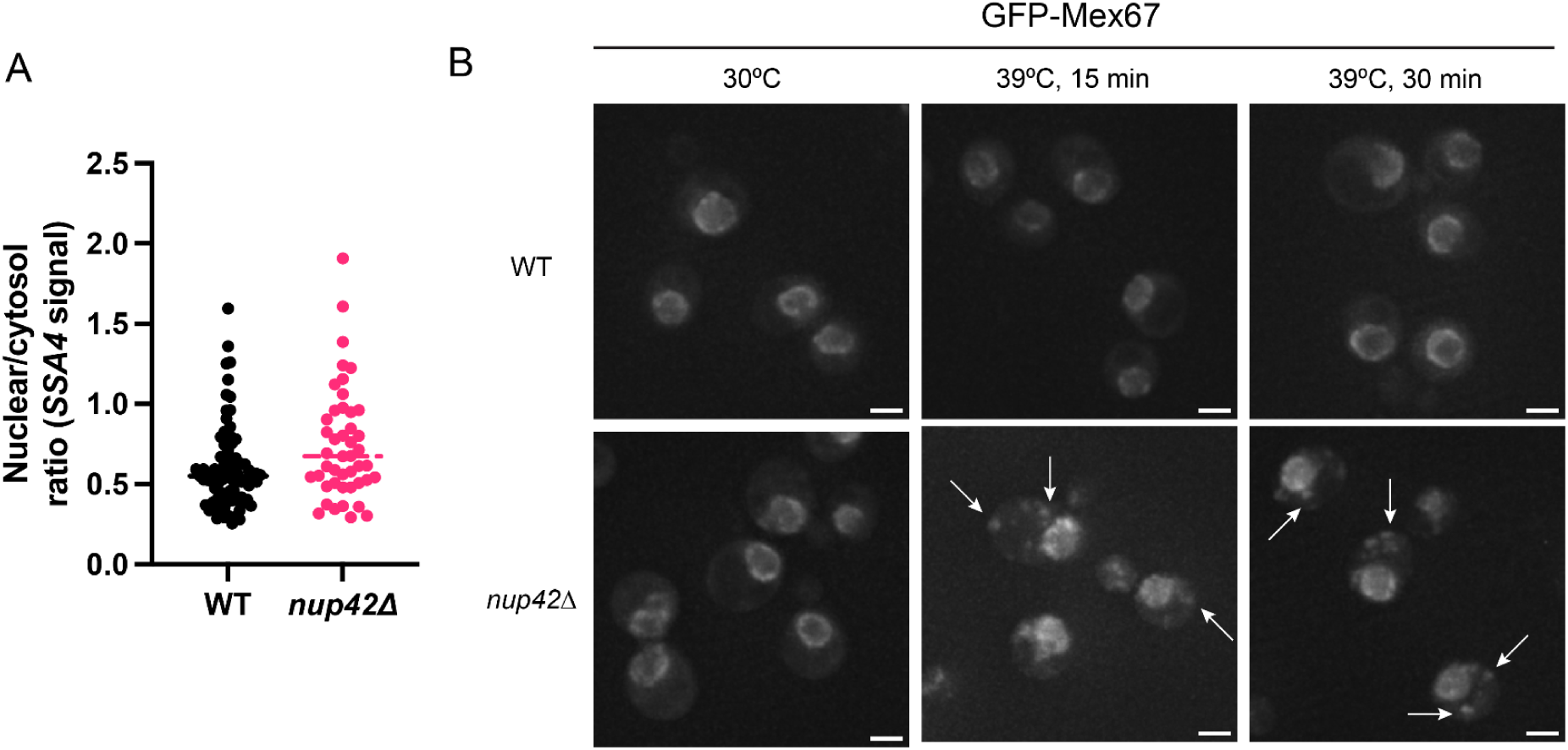
Accumulation of cytosolic Mex67 foci in *nup42Δ* at 39°C. A. Ratiometric quantification of MCP-NLS-2xGFP in the nucleus relative to cytosol in WT and *nup42Δ* at 42°C, 30 minutes. Quantification of the same images represented in Figure 6B. B. Representative Z-stack images of WT and *nup42Δ* GFP-Mex67 cells at 30°C and 39°C heat shock for 15 and 30 minutes. White arrows indicate cytosolic Mex67 foci. Scale bar = 2 µm.

**Figure S6.**
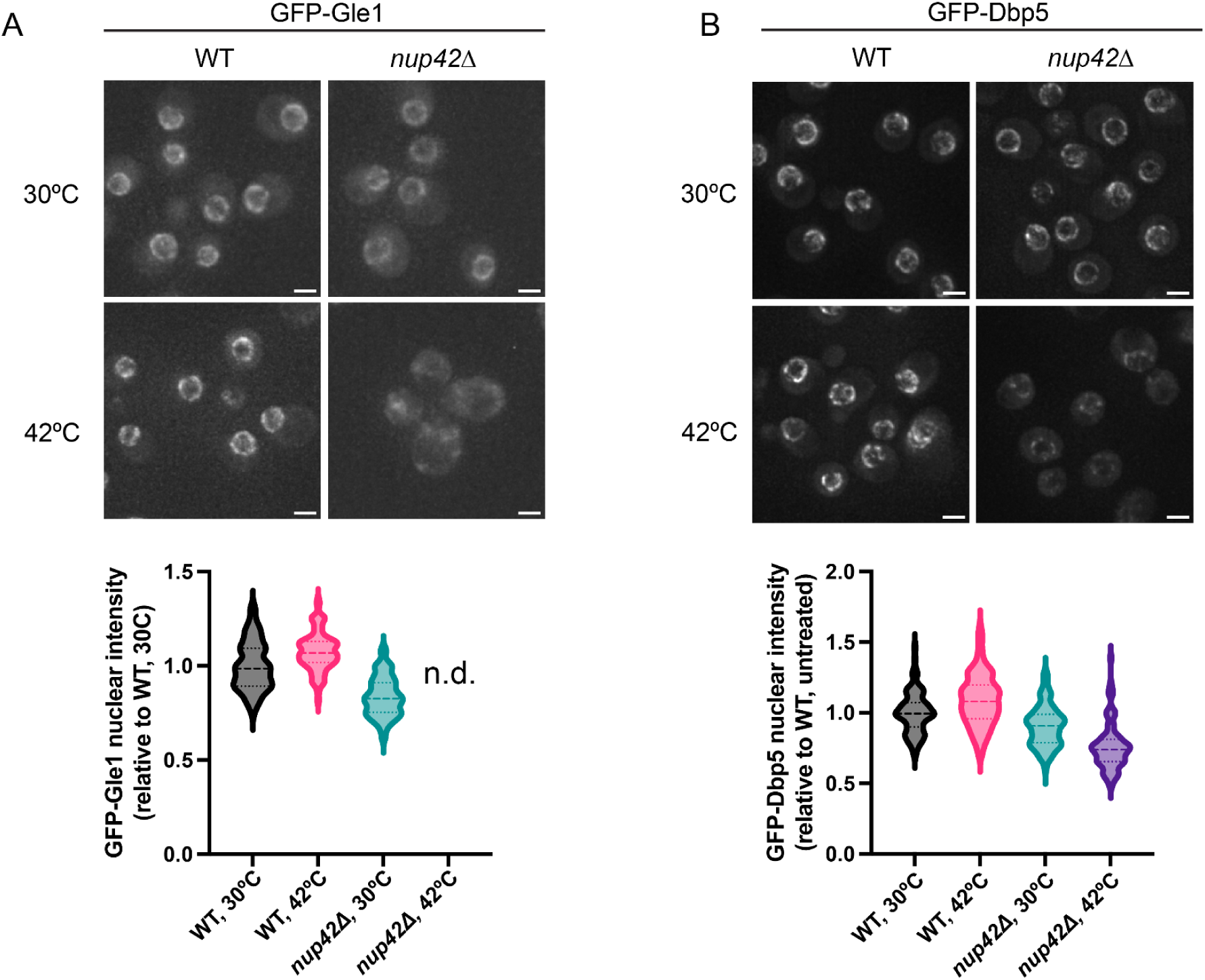
Gle1 and Dbp5 nuclear localization is impaired in the absence of Nup42 during heat shock. A. Top: representative Z-stack images of WT and *nup42Δ* GFP-Gle1 cells at 30°C and 42°C, 30 minutes. Scale bar = 2 µm. Bottom: quantification of GFP-Gle1 nuclear intensity of at least 100 individual cells. n.d. = not detected. B. Top: representative Z-stack images of WT and *nup42Δ* GFP-Dbp5 cells at 30°C and 42°C, 30 minutes. Scale bar = 2 µm. Bottom: quantification of GFP-Dbp5 nuclear intensity of at least 100 individual cells.

**Figure S7.**
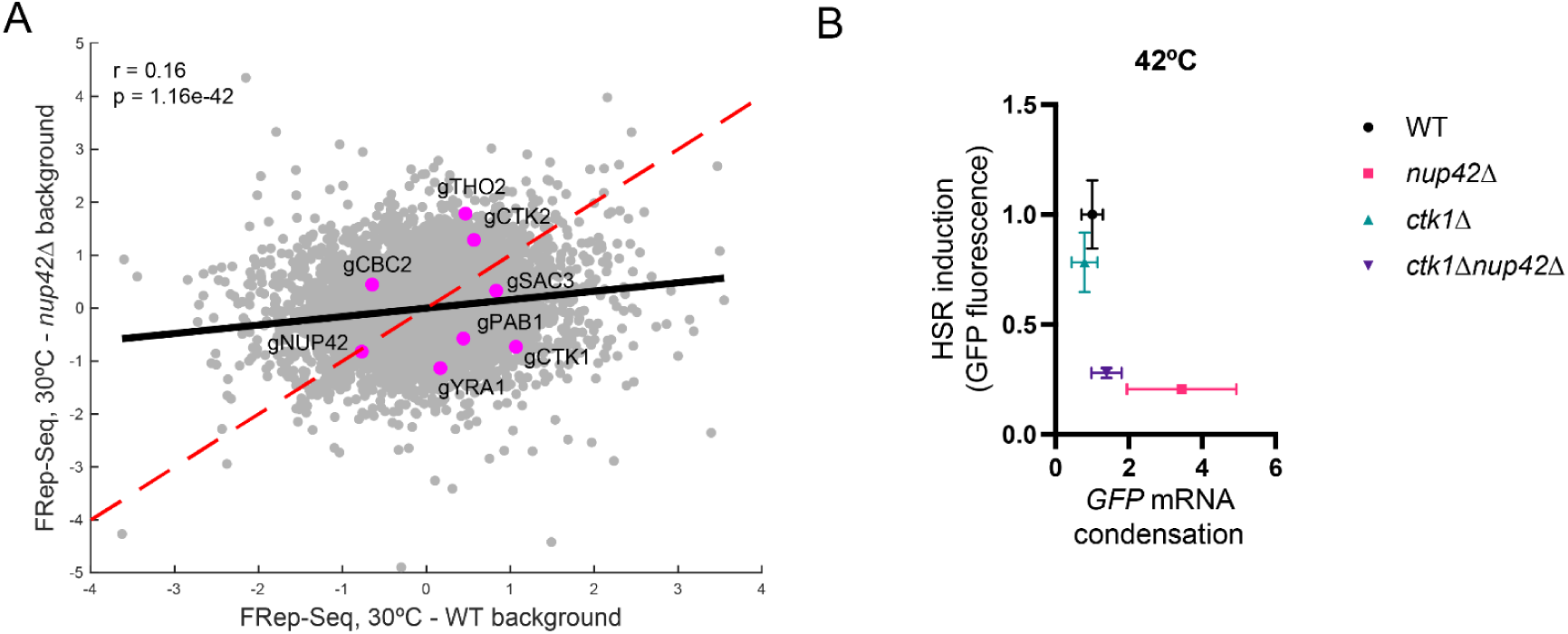
Co-transcriptional mRNP compaction state has minimal effects in mRNP condensation under basal conditions but is a critical determinant in the absence of Nup42 upon heat shock. A. Scatterplot of FRep-Seq screens conducted in WT versus *nup42Δ* background under basal conditions. B. Hsf1-responsive reporter transcript condensation versus protein levels upon 42°C heat shock for 30 minutes, corresponding to data shown in panels 7D and 7E.

## References

1. Warner, J. R. The economics of ribosome biosynthesis in yeast. Trends Biochem. Sci. 24, 437–440 (1999).

2. Lempiäinen, H. & Shore, D. Growth control and ribosome biogenesis. Curr. Opin. Cell Biol. 21, 855–863 (2009).

3. Gasch, A. P. et al. Genomic Expression Programs in the Response of Yeast Cells to Environmental Changes□D.

4. Mühlhofer, M. et al. The Heat Shock Response in Yeast Maintains Protein Homeostasis by Chaperoning and Replenishing Proteins. Cell Rep. 29, 4593–4607.e8 (2019).

5. Albert, B. et al. A ribosome assembly stress response regulates transcription to maintain proteome homeostasis. eLife 8, e45002 (2019).

6. Tye, B. W. et al. Proteotoxicity from aberrant ribosome biogenesis compromises cell fitness. eLife 8, e43002 (2019).

7. Ali, A. et al. Adaptive preservation of orphan ribosomal proteins in chaperone-dispersed condensates. Nat. Cell Biol. 25, 1691–1703 (2023).

8. Ritossa, F. A new puffing pattern induced by temperature shock and DNP in drosophila. Experientia 18, 571–573 (1962).

9. Lindquist, S. THE HEAT-SHOCK RESPONSE. Annu. Rev. Biochem. 55, 1151–1191 (1986).

10. Richter, K., Haslbeck, M. & Buchner, J. The Heat Shock Response: Life on the Verge of Death. Mol. Cell 40, 253–266 (2010).

11. DiDomenico, B. J., Bugaisky, G. E. & Lindquist, S. The heat shock response is self-regulated at both the transcriptional and posttranscriptional levels. Cell 31, 593–603 (1982).

12. Abravaya, K., Phillips, B. & Morimoto, R. I. Attenuation of the heat shock response in HeLa cells is mediated by the release of bound heat shock transcription factor and is modulated by changes in growth and in heat shock temperatures. Genes Dev. 5, 2117–2127 (1991).

13. Baler, R., Zou, J. & Voellmy, R. Evidence for a Role of Hsp70 in the Regulation of the Heat Shock Response in Mammalian Cells. Cell Stress Chaperones 1, 33–39 (1996).

14. Dea, A. & Pincus, D. The Heat Shock Response as a Condensate Cascade. J. Mol. Biol. 436, 168642 (2024).

15. Neef, D. W., Jaeger, A. M. & Thiele, D. J. Heat shock transcription factor 1 as a therapeutic target in neurodegenerative diseases. Nat. Rev. Drug Discov. 10, 930–944 (2011).

16. Hipp, M. S., Kasturi, P. & Hartl, F. U. The proteostasis network and its decline in ageing. Nat. Rev. Mol. Cell Biol. 20, 421–435 (2019).

17. Gomez-Pastor, R., Burchfiel, E. T. & Thiele, D. J. Regulation of heat shock transcription factors and their roles in physiology and disease. Nat. Rev. Mol. Cell Biol. 19, 4–19 (2018).

18. Banani, S. F., Lee, H. O., Hyman, A. A. & Rosen, M. K. Biomolecular condensates: organizers of cellular biochemistry. Nat. Rev. Mol. Cell Biol. 18, 285–298 (2017).

19. Glauninger, H., Wong Hickernell, C. J., Bard, J. A. M. & Drummond, D. A. Stressful steps: Progress and challenges in understanding stress-induced mRNA condensation and accumulation in stress granules. Mol. Cell 82, 2544–2556 (2022).

20. Anderson, P. & Kedersha, N. RNA granules: post-transcriptional and epigenetic modulators of gene expression. Nat. Rev. Mol. Cell Biol. 10, 430–436 (2009).

21. Zid, B. M. & O’Shea, E. K. Promoter sequences direct cytoplasmic localization and translation of mRNAs during starvation in yeast. Nature 514, 117–121 (2014).

22. Iserman, C. et al. Condensation of Ded1p Promotes a Translational Switch from Housekeeping to Stress Protein Production. Cell 181, 818–831.e19 (2020).

23. Desroches Altamirano, C., et al. eIF4F is a thermo-sensing regulatory node in the translational heat shock response. Mol. Cell 84, 1727–1741.e12 (2024).

24. Glauninger, H. et al. Transcriptome-wide mRNP condensation precedes stress granule formation and excludes new mRNAs. Mol. Cell 85, 4393–4409.e11 (2025).

25. Zedan, M. et al. Timing of transcription controls the selective translation of newly synthesized mRNAs during acute environmental stress. Mol. Cell 85, 4379–4392.e5 (2025).

26. Nover, L., Scharf, K.-D. & Neumann, D. Cytoplasmic Heat Shock Granules Are Formed from Precursor Particles and Are Associated with a Specific Set of mRNAs. Mol. Cell. Biol. 9, 1298–1308 (1989).

27. Stöhr, N. et al. ZBP1 regulates mRNA stability during cellular stress. J. Cell Biol. 175, 527–534 (2006).

28. Meinel, D. M. & Sträßer, K. Co-transcriptional mRNP formation is coordinated within a molecular mRNP packaging station in S. cerevisiae. Bioessays 37, 666–677 (2015).

29. Faraway, R., Zenklusen, D. & Plaschka, C. Mechanisms of Messenger RNA Packaging and Export. Annu. Rev. Cell Dev. Biol. 41, 479–504 (2025).

30. Montpetit, B. et al. A conserved mechanism of DEAD-box ATPase activation by nucleoporins and InsP6 in mRNA export. Nature 472, 238–242 (2011).

31. Lin, D. H. et al. Structural and functional analysis of mRNA export regulation by the nuclear pore complex. Nat. Commun. 9, 2319 (2018).

32. Folkmann, A. W., Noble, K. N., Cole, C. N. & Wente, S. R. Dbp5, Gle1-IP6 and Nup159: A working model for mRNP export. Nucleus 2, 540–548 (2011).

33. Seidler, J. F. & Sträßer, K. Understanding nuclear mRNA export: Survival under stress. Mol. Cell 84, 3681–3691 (2024).

34. Asada, R., Dominguez, A. & Montpetit, B. Single-molecule quantitation of RNA-binding protein occupancy and stoichiometry defines a role for Yra1 (Aly/REF) in nuclear mRNP organization. Cell Rep. 42, 113415 (2023).

35. Zander, G. et al. mRNA quality control is bypassed for immediate export of stress-responsive transcripts. Nature 540, 593–596 (2016).

36. Chowdhary, S., Kainth, A. S., Pincus, D. & Gross, D. S. Heat Shock Factor 1 Drives Intergenic Association of Its Target Gene Loci upon Heat Shock. Cell Rep. 26, 18–28.e5 (2019).

37. Chowdhary, S., Kainth, A. S., Paracha, S., Gross, D. S. & Pincus, D. Inducible transcriptional condensates drive 3D genome reorganization in the heat shock response. Mol. Cell 82, 4386–4399.e7 (2022).

38. Chowdhary, S., Paracha, S., Dyer, L. & Pincus, D. Emergent 3D genome reorganization from the stepwise assembly of transcriptional condensates. Preprint at 10.1101/2025.02.23.639564 (2025).

39. Brandman, O. et al. A Ribosome-Bound Quality Control Complex Triggers Degradation of Nascent Peptides and Signals Translation Stress. Cell 151, 1042–1054 (2012).

40. Alford, B. D., et al. ReporterSeq reveals genome-wide dynamic modulators of the heat shock response across diverse stressors. eLife 10, e57376 (2021).

41. Love, M. I., Huber, W. & Anders, S. Moderated estimation of fold change and dispersion for RNA-seq data with DESeq2. Genome Biol. 15, 550 (2014).

42. Pincus, D. et al. Genetic and epigenetic determinants establish a continuum of Hsf1 occupancy and activity across the yeast genome. Mol. Biol. Cell 29, 3168–3182 (2018).

43. Solís, E. J. et al. Defining the Essential Function of Yeast Hsf1 Reveals a Compact Transcriptional Program for Maintaining Eukaryotic Proteostasis. Mol. Cell 63, 60–71 (2016).

44. Teixeira, D. & Parker, R. Analysis of P-Body Assembly in *Saccharomyces cerevisiae*. Mol. Biol. Cell 18, 2274–2287 (2007).

45. Xie, Y. & Varshavsky, A. RPN4 is a ligand, substrate, and transcriptional regulator of the 26S proteasome: A negative feedback circuit. Proc. Natl. Acad. Sci. 98, 3056–3061 (2001).

46. Work, J. J. et al. ReporterSeq reveals genome-wide determinants of proteasome expression. 2021.08.19.456712 Preprint at 10.1101/2021.08.19.456712 (2021).

47. Persson, L. B., Ambati, V. S. & Brandman, O. Cellular Control of Viscosity Counters Changes in Temperature and Energy Availability. Cell 183, 1572–1585.e16 (2020).

48. Kroschwald, S. et al. Promiscuous interactions and protein disaggregases determine the material state of stress-inducible RNP granules. eLife 4, e06807 (2015).

49. Yoo, H., Bard, J. A. M., Pilipenko, E. V. & Drummond, D. A. Chaperones directly and efficiently disperse stress-triggered biomolecular condensates. Mol. Cell 82, 741–755.e11 (2022).

50. Stutz, F. et al. The yeast nucleoporin Rip1p contributes to multiple export pathways with no essential role for its FG-repeat region. Genes Dev. 11, 2857–2868 (1997).

51. Saavedra, C. A., Hammell, C. M., Heath, C. V. & Cole, C. N. Yeast heat shock mRNAs are exported through a distinct pathway defined by Rip1p. Genes Dev. 11, 2845–2856 (1997).

52. Vainberg, I. E., Dower, K. & Rosbash, M. Nuclear Export of Heat Shock and Non-Heat-Shock mRNA Occurs via Similar Pathways. Mol. Cell. Biol. 20, 3996–4005 (2000).

53. Adams, R. L., Terry, L. J. & Wente, S. R. Nucleoporin FG Domains Facilitate mRNP Remodeling at the Cytoplasmic Face of the Nuclear Pore Complex. Genetics 197, 1213–1224 (2014).

54. Adams, R. L., Mason, A. C., Glass, L., Aditi & Wente, S. R. Nup42 and IP_6_ coordinate Gle1 stimulation of Dbp5/_DDX19B for MRNA export in yeast and human cells. Traffic 18, 776–790 (2017).

55. Segref, A. et al. Mex67p, a novel factor for nuclear mRNA export, binds to both poly(A)+ RNA and nuclear pores. EMBO J. 16, 3256–3271 (1997).

56. Snay-Hodge, C. A., Colot, H. V., Goldstein, A. L. & Cole, C. N. Dbp5p/Rat8p is a yeast nuclear pore-associated DEAD-box protein essential for RNA export. EMBO J. 17, 2663–2676 (1998).

57. Gorsch, L. C., Dockendorff, T. C. & Cole, C. N. A conditional allele of the novel repeat-containing yeast nucleoporin RAT7/NUP159 causes both rapid cessation of mRNA export and reversible clustering of nuclear pore complexes. J. Cell Biol. 129, 939–955 (1995).

58. Murphy, R. & Wente, S. R. An RNA-export mediator with an essential nuclear export signal. Nature 383, 357–360 (1996).

59. Jensen, T. H., Patricio, K., McCarthy, T. & Rosbash, M. A Block to mRNA Nuclear Export in S. cerevisiae Leads to Hyperadenylation of Transcripts that Accumulate at the Site of Transcription. Mol. Cell 7, 887–898 (2001).

60. Tutucci, E. et al. An improved MS2 system for accurate reporting of the mRNA life cycle. Nat. Methods 15, 81–89 (2018).

61. Bonneau, F. et al. Nuclear mRNPs are compact particles packaged with a network of proteins promoting RNA–RNA interactions. Genes Dev. 37, 505–517 (2023).

62. Fischer, T. et al. The mRNA export machinery requires the novel Sac3p–Thp1p complex to dock at the nucleoplasmic entrance of the nuclear pores. EMBO J. 21, 5843–5852 (2002).

63. Köhler, A. & Hurt, E. Exporting RNA from the nucleus to the cytoplasm. Nat. Rev. Mol. Cell Biol. 8, 761–773 (2007).

64. Jani, D. et al. Functional and structural characterization of the mammalian TREX-2 complex that links transcription with nuclear messenger RNA export. Nucleic Acids Res. 40, 4562–4573 (2012).

65. Ahn, S. H., Keogh, M. & Buratowski, S. Ctk1 promotes dissociation of basal transcription factors from elongating RNA polymerase II. EMBO J. 28, 205–212 (2009).

66. Meinel, D. M. et al. Recruitment of TREX to the Transcription Machinery by Its Direct Binding to the Phospho-CTD of RNA Polymerase II. PLOS Genet. 9, e1003914 (2013).

67. Qiu, H., Hu, C. & Hinnebusch, A. G. Phosphorylation of the Pol II CTD by KIN28 Enhances BUR1/BUR2 Recruitment and Ser2 CTD Phosphorylation Near Promoters. Mol. Cell 33, 752–762 (2009).

68. Xie, Y. et al. Structure and activation mechanism of the yeast RNA Pol II CTD kinase CTDK-1 complex. Proc. Natl. Acad. Sci. 118, e2019163118 (2021).

69. Strahm, Y. The RNA export factor Gle1p is located on the cytoplasmic fibrils of the NPC and physically interacts with the FG-nucleoporin Rip1p, the DEAD-box protein Rat8p/Dbp5p and a new protein Ymr255p. EMBO J. 18, 5761–5777 (1999).

70. Van Treeck, B. et al. RNA self-assembly contributes to stress granule formation and defining the stress granule transcriptome. Proc. Natl. Acad. Sci. 115, 2734–2739 (2018).

71. Van Treeck, B. & Parker, R. Emerging Roles for Intermolecular RNA-RNA Interactions in RNP Assemblies. Cell 174, 791–802 (2018).

72. Khong, A. & Parker, R. The landscape of eukaryotic mRNPs. RNA 26, 229–239 (2020).

73. Navalkar, A., Eppert, M., Sabari, B. R. & Mittag, T. Density transitions in the regulation of transcription. Mol. Cell 0, (2026).

74. Carmody, S. R., Tran, E. J., Apponi, L. H., Corbett, A. H. & Wente, S. R. The Mitogen-Activated Protein Kinase Slt2 Regulates Nuclear Retention of Non-Heat Shock mRNAs during Heat Shock-Induced Stress. Mol. Cell. Biol. 30, 5168–5179 (2010).

75. Dekker, M., Van Der Giessen, E. & Onck, P. R. Phase separation of intrinsically disordered FG-Nups is driven by highly dynamic FG motifs. Proc. Natl. Acad. Sci. 120, e2221804120 (2023).

76. Patel, S. S., Belmont, B. J., Sante, J. M. & Rexach, M. F. Natively Unfolded Nucleoporins Gate Protein Diffusion across the Nuclear Pore Complex. Cell 129, 83–96 (2007).

77. Krueger, F., James, F., Ewels, P., Afyounian, E. & Schuster-Boeckler, B. FelixKrueger/TrimGalore: v0.6.7 - DOI via Zenodo. Zenodo 10.5281/zenodo.5127899 (2021).

78. Dobin, A. et al. STAR: ultrafast universal RNA-seq aligner. Bioinformatics 29, 15–21 (2013).

79. Yu, G., Wang, L.-G., Han, Y. & He, Q.-Y. clusterProfiler: an R package for comparing biological themes among gene clusters. Omics J. Integr. Biol. 16, 284–287 (2012).

80. Fang, F. et al. A vector set for systematic metabolic engineering in Saccharomyces cerevisiae. Yeast Chichester Engl. 28, 123–136 (2011).

